# The Arabidopsis MCTP member QUIRKY regulates the formation of the STRUBBELIG receptor kinase complex

**DOI:** 10.1101/2022.09.20.508642

**Authors:** Xia Chen, Barbara Leśniewska, Prasad Vaddepalli, Kay Schneitz

## Abstract

Intercellular communication plays a central role in organogenesis. Tissue morphogenesis in *Arabidopsis thaliana* requires signaling mediated by a cell surface complex containing the atypical receptor kinase STRUBBELIG (SUB) and the multiple C2 domains and transmembrane region protein QUIRKY (QKY). QKY is required to stabilize SUB at the plasma membrane. However, it is unclear what the in vivo architecture of the QKY/SUB signaling complex is, how it is controlled, and how it relates to the maintenance of SUB at the cell surface. Using a combination of yeast two-hybrid assays and Förster resonance energy transfer (FRET)/fluorescence lifetime imaging microscopy (FLIM) in epidermal cells of seedling roots we find that QKY promotes the formation of SUB homo-oligomers in vivo, a process that appears to involve an interaction between the extracellular domains of SUB. We also show that QKY and SUB physically interact and form a complex at the cell surface in vivo. In addition, the data show that the N-terminal C2A-B region of QKY interacts with the intracellular domain of SUB. They further reveal that this interaction is essential to maintain SUB levels at the cell surface. Finally, we provide evidence that QKY forms homo-multimers in vivo in a SUB-independent manner. We suggest a model in which the physical interaction of QKY with SUB mediates the oligomerization of SUB and attenuates its internalization, thereby maintaining sufficiently high levels of SUB at the cell surface required for the control of tissue morphogenesis.

## Introduction

Tissue morphogenesis is a complex process that depends on sophisticated cell-to-cell communication. Receptor kinases (RKs) represent central components of cell-surface receptor complexes involved in extracellular ligand perception in plants (Hohmann *et al*, 2017). In Arabidopsis, the atypical RK STRUBBELIG (SUB) (Chevalier *et al*, 2005), also known as SCRAMBLED (SCM) (Kwak *et al*, 2005), participates in multiple developmental and stress pathways. SUB controls root hair patterning, leaf development, floral organ shape, and ovule development (Chevalier *et al*, 2005; Kwak *et al*, 2005; Fulton *et al*, 2009; Lin *et al*, 2012). Recently, SUB has also been shown to be required for the stress response elicited by a reduction in cellulose biosynthesis (Chaudhary *et al*, 2020, 2021). Genetic evidence suggested that SUB functions as a scaffold protein in several cell surface signaling complexes (Vaddepalli *et al*, 2011; Chaudhary *et al*, 2021). In support of this notion SUB harbors an ECD with only six LRRs (Vaddepalli *et al*, 2011) and thus is akin to the ECD of known co-receptors, such as BRI1-ASSOCIATED RECEPTOR KINASE 1/SOMATIC EMBROYGENESIS RECEPTOR-LIKE KINASE 3 (BAK1/SERK3) that carries five LRRs (Li *et al*, 2002; Nam & Li, 2002). BAK1 and other SERK members are implied in the control of growth, development, and immunity by acting as co-receptors for several different signaling RKs, including BRASSINOSTEROID-INSENSITIVE 1 (BRI1) and FLAGELLIN SENSITIVE 2 (FLS2) (Chinchilla *et al*, 2007; She *et al*, 2011; Hothorn *et al*, 2011; Liu *et al*, 2020).

A signaling RK with which SUB partners has yet to be identified. QUIRKY (QKY) represents the only known other factor of a SUB signaling complex in vivo. QKY is essential for SUB signaling involved in tissue morphogenesis but plays a minor role in the cell wall stress response (Trehin *et al*, 2013; Fulton *et al*, 2009; Song *et al*, 2019; Chaudhary *et al*, 2020). Several lines of evidence suggest that QKY and SUB are components of a complex (Trehin *et al*, 2013; Vaddepalli *et al*, 2014; Song *et al*, 2019) and physically interact at plasmodesmata (PD) in epidermal root cells of young Arabidopsis seedlings (Vaddepalli *et al*, 2014). SUB undergoes constitutive clathrin-dependent endocytosis followed by degradation in the vacuole (Gao *et al*, 2019; Song *et al*, 2019) and QKY is also required for the stabilization of SUB at the cell surface (Song *et al*, 2019; Chaudhary *et al*, 2021). Apart from its function in SUB-dependent signaling, QKY has been reported to play other roles in growth and development.Genetic evidence indicates that *QKY* and its maize homolog *Carbohydrate Partitioning Defective33* (*cpd33*) play a role in sugar export from leaves and thus carbohydrate partitioning (Tran *et al*, 2019). In addition, *QKY* participates in the control of flowering time (Liu *et al*, 2019), a process in which SUB does not seem to play a role.

QKY belongs to the family of multiple C2 domains and transmembrane region proteins (MCTPs). MCTPs carry three to four lipid-binding C2 domains in the N-terminal half and multiple transmembrane domains in the C-terminal region (Shin *et al*, 2005). Representatives of the plant MCTP family were identified in algae and numerous land plant species (Fulton *et al*, 2009; Liu *et al*, 2012; Zhu *et al*, 2020; Liu *et al*, 2018a; Brault *et al*, 2019; Tran *et al*, 2019; Hao *et al*, 2020; Hu *et al*, 2021).Several studies indicated multiple members of the MCTP family to be present at PD (Fernandez-Calvino *et al*, 2011; Liu *et al*, 2012; Vaddepalli *et al*, 2014; Tran *et al*, 2019; Brault *et al*, 2019). PD are considered plant-specific membrane contact sites (MCS) between the endoplasmic reticulum (ER) and the plasma membrane (PM) (Tilsner *et al*, 2016). The structure of MCTPs and their presence at PD led to the proposal that some MCTPs function as plasmodesmal membrane tethers (Brault *et al*, 2019). In addition to QKY, the function of several other plant MCTPs has been described. MCTP1/FT-INTERACTING PROTEIN 1 (FTIP1) controls flowering time in Arabidopsis and rice (Liu *et al*, 2012; Song *et al*, 2017), while MCTP3 and MCTP4 affect meristem development (Liu *et al*, 2018b; Brault *et al*, 2019), as do GhMCTP7, GhMCTP12, and GhMCTP17 in cotton (Hu *et al*, 2021).

Many aspects of the QKY/SUB signaling complex in its native cellular environment are unknown. In this study, we provide an in-depth description of the QKY subcellular localization. We further tested the ability of SUB and QKY to form multimers. We mapped the SUB interaction domain of QKY and analyzed its topology in the SUB/QKY complex. Thus, we provide new molecular insights into the QKY-dependent control of SUB complex architecture in vivo.

## Results

### QKY localizes to the ER and to PD

Experiments involving stable transgenic Arabidopsis plants carrying a functional QKY:EGFP reporter indicated a PD localization for QKY (Vaddepalli *et al*, 2014). Transient expression studies in *Nicotiana benthamiana* leaves frequently suggested ER localization of MCTP proteins, but also indicated a number of different subcellular associations, depending on the family member (Liu *et al*, 2018a; Brault *et al*, 2019; Liu *et al*, 2019; Tran *et al*, 2019; Hu *et al*, 2021). In particular, they suggested that QKY is localized to the ER, intracellular vesicles, PM, and cytoplasm. We reinvestigated the subcellular localization of QKY in its native context in Arabidopsis cells. First, we transformed the *qky-9* mutant, harboring a null allele (Fig. S1), with a reporter carrying an N-terminal translational fusion of mCherry to QKY driven by its endogenous promoter (pQKY::mCherry:QKY) (see Materials and Methods) and analyzed the phenotype of 17 independent transformant lines (Table1). We found that pQKY::mCherry:QKY robustly restored the floral tissue and root hair patterning defects of *qky-9* (Table1, Table 2) (Fig. S2).

**Table 1.** Phenotypes of *qky-9* lines homozygous for different QKYdel variants.

| Genotype | Phenotype<sup>a</sup> | Percentage of independent homozygous T3 lines<sup>b</sup> |
| --- | --- | --- |
| QMQ *qky-9* | WT | 15/17 |
| delC2A *qky-9* | *qky* | 10/10 |
| delC2AL *qky-9* | *qky* | 12/12 |
| delC2A-B *qky-9* | *qky* | 11/11 |
| delC2A-C *qky-9* | *qky* | 6/6 |
| delC2A-D *qky-9* | *qky* | 9/9 |

<sup>a</sup>Phenotype was scored based on the morphology of flowers, siliques, and stems.
<sup>b</sup>Number of independent transgenic lines with corresponding phenotype / total number of independent transgenic lines scored. Abbreviations: QMQ, pQKY::mCherry:QKY.

**Table 2.** Integument defects of L*er, qky-9, qky-9 pQKY::mCherry:QKY*, and different *qky-9 pQKY::mCherry:QKY*del lines.

| Genotype | N ovules scored<sup>a</sup> | % defective ovules<sup>b</sup> |
| --- | --- | --- |
| *Ler* | 324 | 3.40 |
| *qky-9* | 428 | 36.92 |
| QMQ *qky-9* | 396 | 4.55 |
| delC2A *qky-9* | 371 | 34.23 |
| delC2AL *qky-9* | 422 | 33.89 |
| delC2A-B *qky-9* | 463 | 32.83 |
| delC2A-C *qky-9* | 367 | 35.15 |
| delC2A-D *qky-9* | 335 | 39.70 |

For *Ler* and *qky-9*, 7 to 10 different carpels from at least two different plants were analyzed. For each *qky-9* pQKY::mCherry:QKYdel genotype, 7 to 10 different carpels from at least two different plants of two independent lines each were analyzed. Abbreviations: QMQ, pQKY::mCherry:QKY.
<sup>a</sup>Total number of ovules scored.
<sup>b</sup>Percentage of ovules with *qky*-like integument defects.

Second, we analyzed the subcellular localization of the pQKY::mCherry:QKY reporter in six of these lines in live tissues by confocal laser scanning microscopy (CLSM). We observed a modest signal with undistinguishable subcellular signal distribution between all six lines. For further analysis we used line 6 which exhibited average signal intensity. We detected reporter signals in several tissues, including the root epidermis of 6-day seedlings and several stage 13 floral organs (Fig. 1) (floral stages according to (Smyth *et al*, 1990)). As previously reported, we observed a spotty PD-related pattern along the cell periphery of the root epidermis of seedlings (Fig. 1A-C) (Vaddepalli *et al*, 2014). In addition, we detected a faint ER-like signal. A similar pattern, though with a somewhat less prominent ER-like and PD-related signal, was observed in the epidermis of mature sepals, petals, and carpels (Fig. 1D-L). An endosomal localization of a p35S::GFP:QKY reporter was observed after transient expression in tobacco leaves in line with its suggested function in the vesicular transport of FLOWERING TIME (FT) in leaf phloem companion cells (Liu *et al*, 2019). We never observed intracellular vesicle-like signals in the analyzed cells of our Arabidopsis lines with our reporters. Third, we performed a co-localization experiment. As a control, we stained lateral root cap cells of 6-day seedlings expressing Q4, a well-established marker for the ER (Cutler *et al*, 2000; Cutler & Ehrhardt, 2002), with the membrane marker FM4-64 for 5 minutes, thereby labeling the PM, and investigated the subcellular distribution of Q4 and FM4-64-derived signals. We detected very little if any overlap between two signals in these cells (Fig. 1M-Q). We then crossed Q4 and pQKY::mCherry:QKY lines and analyzed the subcellular distribution of the two reporters in the same cells of F1 6-day seedlings. pQKY::mCherry:QKY localization in lateral root cap cells exhibited a prominent ER-like pattern (Fig. 1Q,R) and we could readily observe co-localization of Q4 and pQKY::mCherry:QKY signals in these cells (Fig. 1R-V). In summary, our data are compatible with the notion that QKY localizes to the ER and PD in Arabidopsis cells.

**Figure 1:**
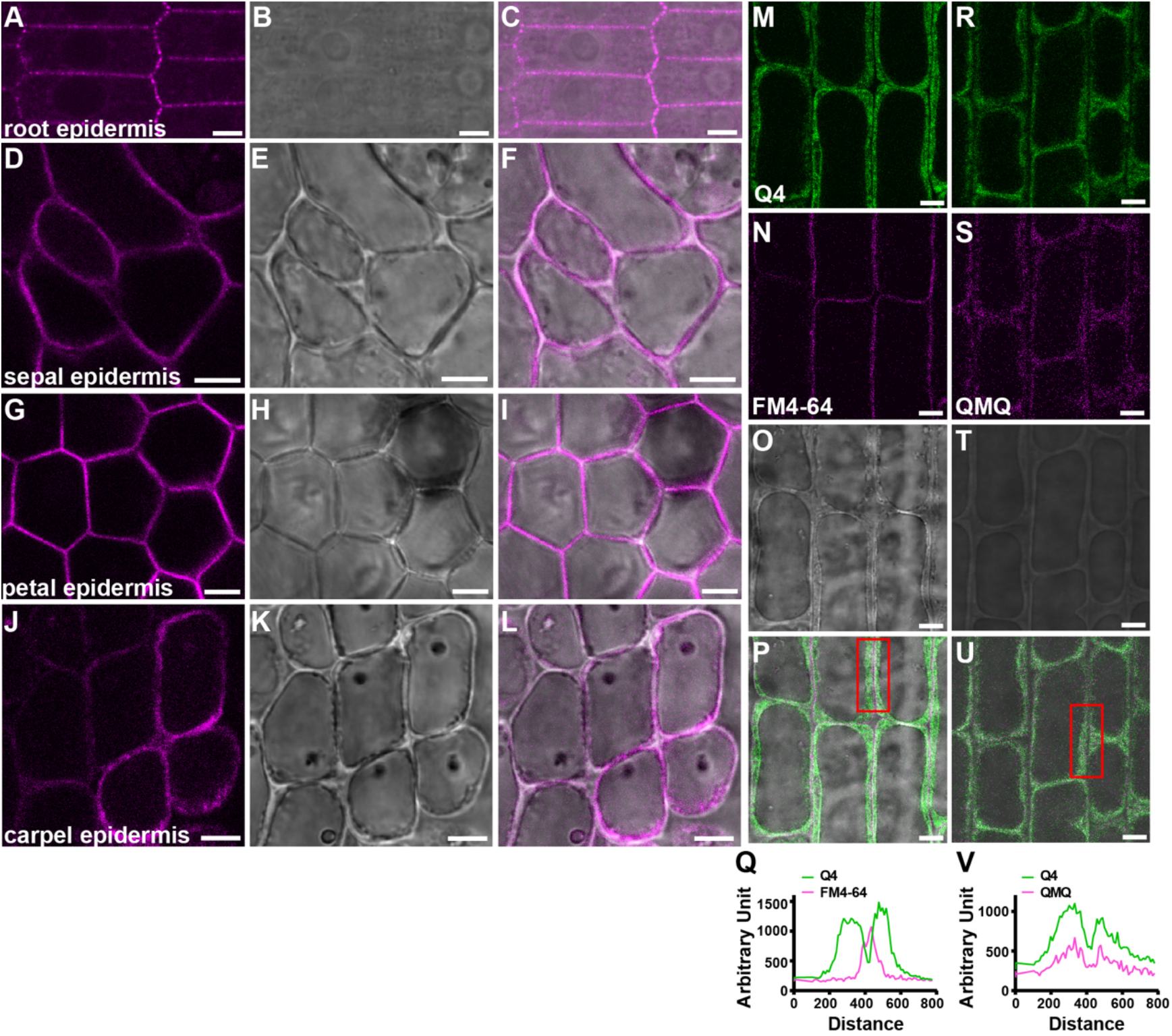
Subcellular localization of the pQKY::mCherry:QKY reporter in different tissues of *qky-9*. Confocal micrographs are shown. (A-D) Epidermal cells of different tissues. (A) 6-days-old seedling root. (B-D) Different floral organs at floral stage 13. The floral organs are indicated. Similar arrangement as in (A). (M-Q) Signal distribution of the ER-resident Q4 marker and the PM-labeling FM4-64 stain, respectively, in lateral root cap cells. (M) The GFP-derived signal of the Q4 marker. (N) FM4-64 signal. (O) The DIC channel. (P) Merge of (M-O). (Q) Intensity profiles measured along a line connecting the dots highlighted by the red rectangle in (P). The x-axis marks the arbitrary distance. The y-axis denotes arbitrary intensity units. (R-V) Colocalization of Q4 and mCherry:QKY signals. (R) Q4 signal. (S) mCherry:QKY signal. (T) DIC. (U) Merge of (R-T). (V) Intensity profiles measured along a line connecting the dots highlighted by the red rectangle in (U). The x-axis marks the arbitrary distance. The y-axis denotes arbitrary intensity units.. Abbreviation: QMQ, pQKY::mCherry:QKY *qky-9*. Scale bars: 5 μm.

### QKY undergoes SUB-independent homo-oligomerization

Maize MCTP CPD33 was reported to form homodimers in bimolecular fluorescence complementation (BiFC) assays in tobacco leaf cells (Tran *et al*, 2019). We therefore investigated whether QKY forms homo-multimers. We first performed a yeast two-hybrid (Y2H) experiment in which the region spanning the C2A to C2D domains (C2A-D) served as bait and prey. We observed yeast growth on the selective medium, indicating that this region can interact with itself in this assay (Fig. 2). To map the interaction domain, we generated a series of deletions within the C2A-D domain and tested their ability to interact with the C2A-D domain. The results indicate that the C2A-B domain is necessary for interaction in yeast (Fig. 2).

**Figure 2.**
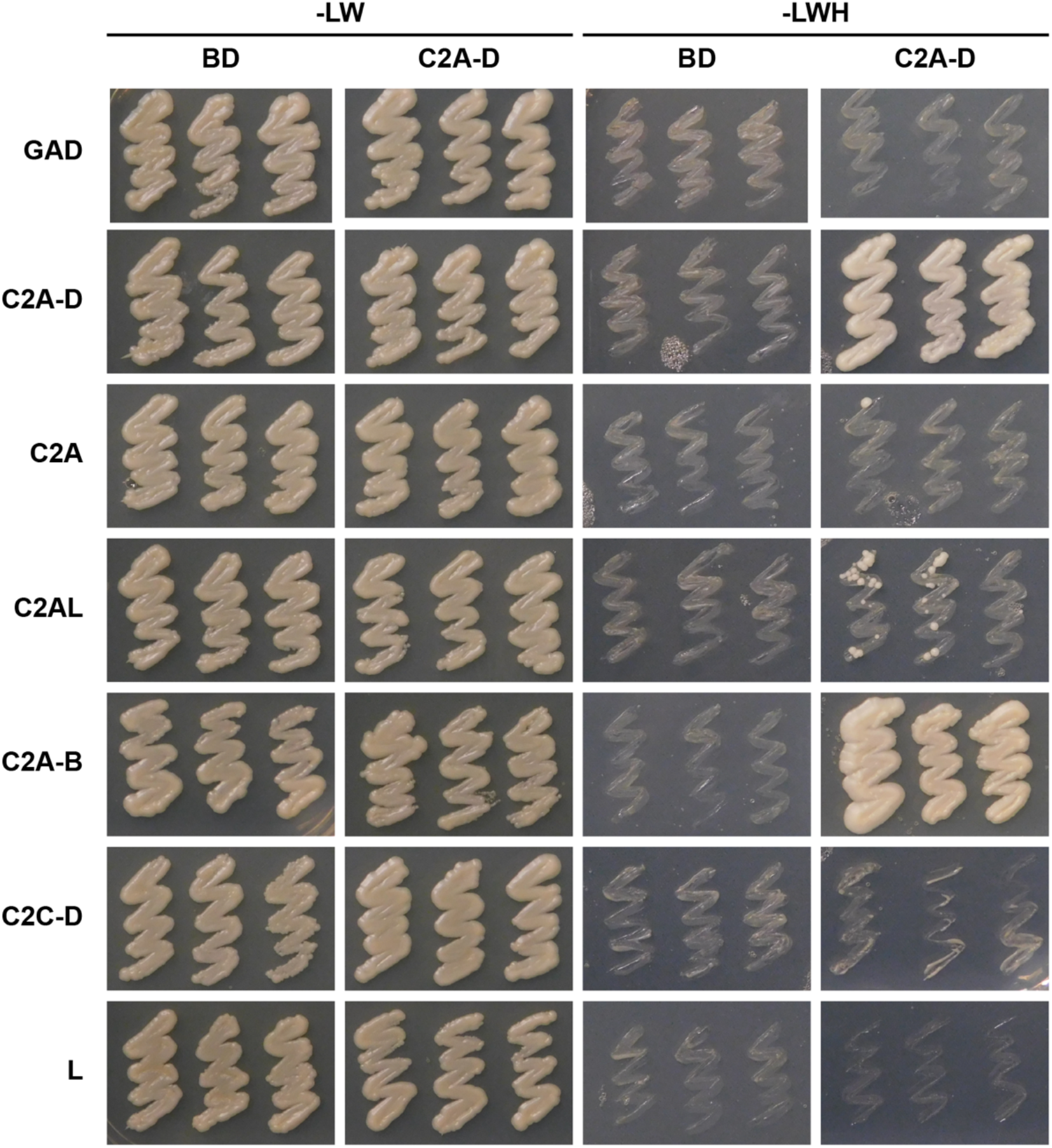
QKY homo-dimerizes in a yeast two-hybrid assay. Yeast two-hybrid assay involving the QKY C2A-D domain or various deletions fused to the GAL4 activating domain (GAD) and QKY C2A-D domain fused to the GAL4 DNA-binding domain (GBD). Growth on -LW panel indicates successful transformation of both plasmids and on -LWH panel indicates presence or absence of interaction. Results from three independent transformation events are shown. The experiment was repeated twice with identical results.

Next, we tested if QKY can form homo-multimers in its native environment. To this end, we tested the interaction in epidermal root cells of 5-day seedlings using steady-state Förster resonance energy transfer (FRET)/fluorescence lifetime imaging microscopy (FRET/FLIM). We analyzed 5-day seedlings expressing functional pUBQ10::EGFP:QKY or pUBQ10::EGFP:QKY and pUBQ10::mCherry:QKY reporters. When we analyzed the subcellular distribution of the PD-related pQKY::mCherry:QKY signals, we detected stronger signals along the longitudinal cell periphery (Fig. S3A) (Vaddepalli *et al*, 2014). Therefore, we distinguished between the transverse PM, the longitudinal PM, and the PD located along the longitudinal PM. We found that the mean fluorescence lifetime of pUBQ10::EGFP:QKY was significantly reduced when UMQ was present indicating that pUBQ10::EGFP:QKY forms homo-oligomers in Arabidopsis cells (Fig. 3A). As negative control, we used a line co-expressing the PM-localized p35S::Lti6a:GFP (Lti6a:GFP) (Cutler *et al*, 2000) and p2×35S::Lti6b:2xmCherry (Lti6b:2xmCherry) (Noack *et al*, 2022) reporters. We never observed significant mean fluorescence lifetime reductions for Lti6a:GFP in the presence of Lti6b:2xmCherry (Fig. 3B). The combined results indicate that QKY, and possibly other MCTP proteins, undergo homo-oligomerization in vivo.

**Figure 3.**
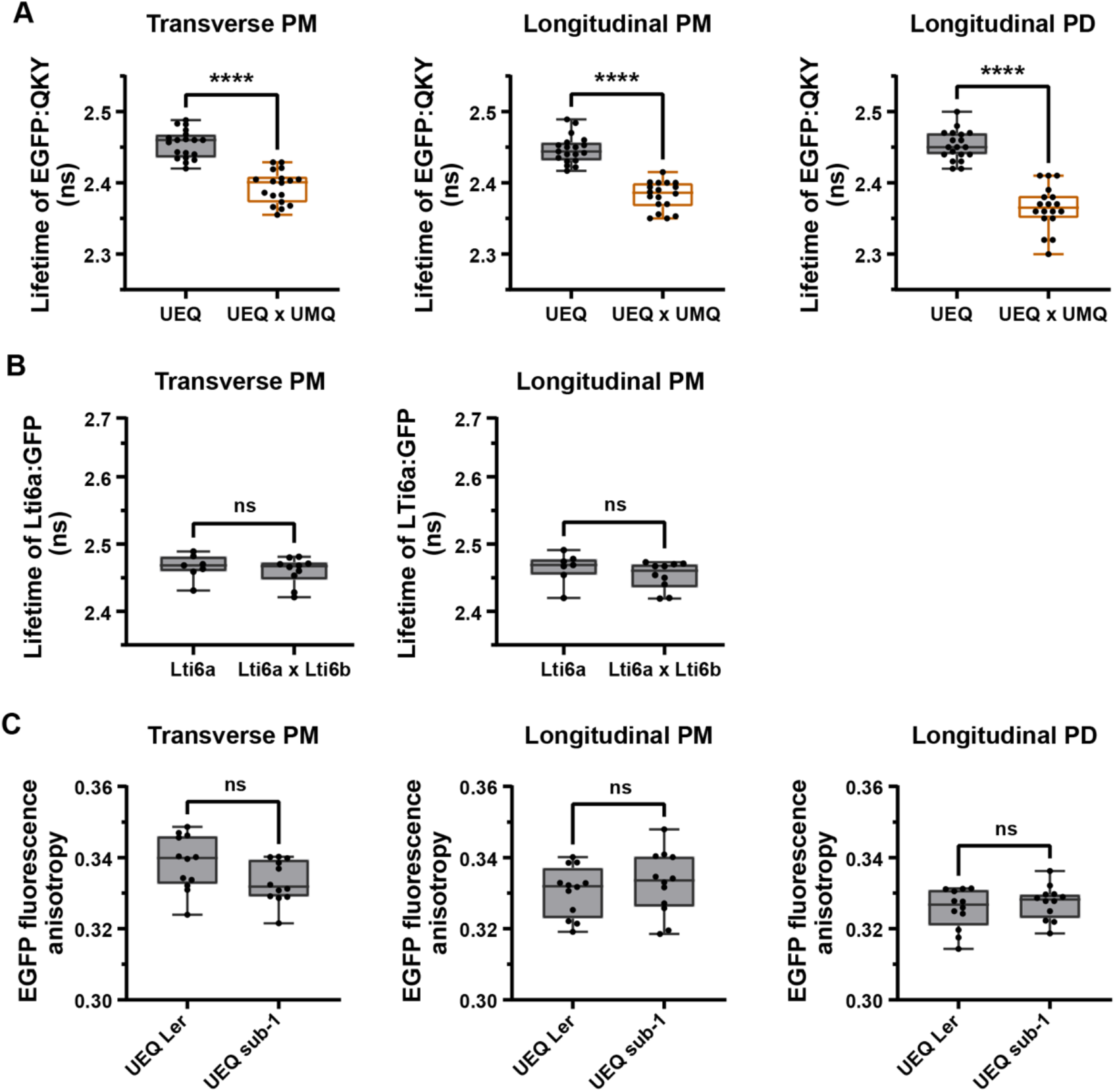
QKY undergoes SUB-independent homo-oligomerization in vivo. (A) FRET-FLIM analysis in stably transgenic plants harboring EGFP:QKY or EGFP:QKY combined with mCherry:QKY at the transverse PM, longitudinal PM and longitudinal PD, respectively. Experiments were in L*er* background. Box and whiskers plots are shown. 18≤n≤19, each data point represents a mean of at least five cells per root. ****P<0.0001. Unpaired t test, two-tailed P-values. (B) Quantification of the Lti6a:GFP fluorescence lifetime in stably transgenic Lti6a:GFP and Lti6a:GFP/Lti6b:2xmCherry plants at the transverse and longitudinal PM, respectively. Box and whisker plots are shown. 7≤n≤10, each data point represents a mean of at least five cells per root. ns: not significant. Unpaired t test, two-tailed P-values. (C) Analysis of EGFP:QKY fluorescence anisotropy in the indicated genetic backgrounds at the transverse PM, longitudinal PM and longitudinal PD, respectively. Box and whisker plots are shown. 12≤n≤13, each data point represents a mean of at least five cells per root. ns: not significant. Unpaired t test, two-tailed P-values. Experiments were repeated at least two times with similar results. Box and whiskers plots show the median values (middle bars) and the 25th to 75th percentiles (box). The whiskers show the smallest and largest values. Abbreviations: CW, cell wall; DM, desmotubule; ER, endoplasmic reticulum; Lti6a, p35S::Lti6a:GFP; Lti6b, p2×35S::Lti6b:2xmCherry; PM, plasma membrane; UEQ, pUBQ10::EGFP:QKY; UMQ, pUBQ10::mCherry:QKY.

We then explored whether QKY oligomerization was dependent on SUB. To this end we measured the steady-state fluorescence anisotropy (FA) of pUBQ10::EGFP:QKY in epidermal root cells of 5-day wild-type and *sub-1* seedlings. Fluorescence anisotropy is a measure of the rotational freedom of a fluorescent molecule, such as GFP. Upon protein homo-oligomerization of GFP-based fusion proteins homo-FRET can occur resulting in a decrease in fluorescence anisotropy (Bader *et al*, 2011; Weidtkamp-Peters & Stahl, 2017) (Fig. S4). This approach has been successfully applied in receptor kinase interaction studies involving for example CLAVATA1 or BAK1 (Somssich *et al*, 2015; Stahl *et al*, 2013). We did not find any significant differences in the FA values of pUBQ10::EGFP:QKY in wild type compared with *sub-1* (Fig. 3C). The combined results support the model that QKY undergoes SUB-independent homo-oligomerization in vivo and that this interaction is mediated by the C2A-B domain.

### QKY physically interacts with SUB in vivo

Previous results suggested the physical interaction of QKY and SUB at PD (Vaddepalli *et al*, 2014). To confirm the QKY/SUB interaction we generated novel lines expressing a newly generated translational fusion of SUB to GFP driven by the endogenous SUB promoter (pSUB::SUB:GFP) and the functional mCherry::QKY fusion driven by the endogenous *QKY* promoter (pQKY::mCherry:QKY). The pSUB::SUB:GFP reporter is functional as it complements the null allele *sub-1* (Fig. S5). Taking advantage of an advanced microscopy set up (see Materials and Methods) we performed FRET/FLIM experiments in root epidermal cells of 5-day seedlings (Fig. 4A-I). As outlined above, we distinguished between the transverse PM, the longitudinal PM, and the PD located along the longitudinal PM. We detected a significant decrease in the mean fluorescent lifetime of pSUB::SUB:GFP in the presence of pQKY::mCherry:QKY. We also observed a polar distribution of the reduction in mean fluorescence lifetime of pSUB::SUB:GFP in the presence of pQKY::mCherry:QKY (Fig. 4I). The reduction in pSUB::SUB:GFP mean fluorescence lifetime was more pronounced at the longitudinal PM than at the transverse PM, consistent with the observed polar distribution of signal intensity of pSUB::SUB:GFP and pQKY::mCherry:QKY. We could not detect an obvious difference in the reduction of pSUB::SUB:GFP mean fluorescence lifetime when we compared the longitudinal PM with the PD-related signals along the longitudinal PM (Fig. 4I). As negative control, we used a line co-expressing pSUB::SUB:GFP and the Lti6b:2xmCherry reporter. We never observed significant mean fluorescence lifetime reductions for pSUB::SUB:GFP in the presence of Lti6b:2xmCherry (Fig. 4I). The results support the notion of a physical interaction between QKY and SUB in vivo. Interestingly, they also suggest that QKY and SUB preferentially interact at the longitudinal edges of epidermal root cells.

**Figure 4.**
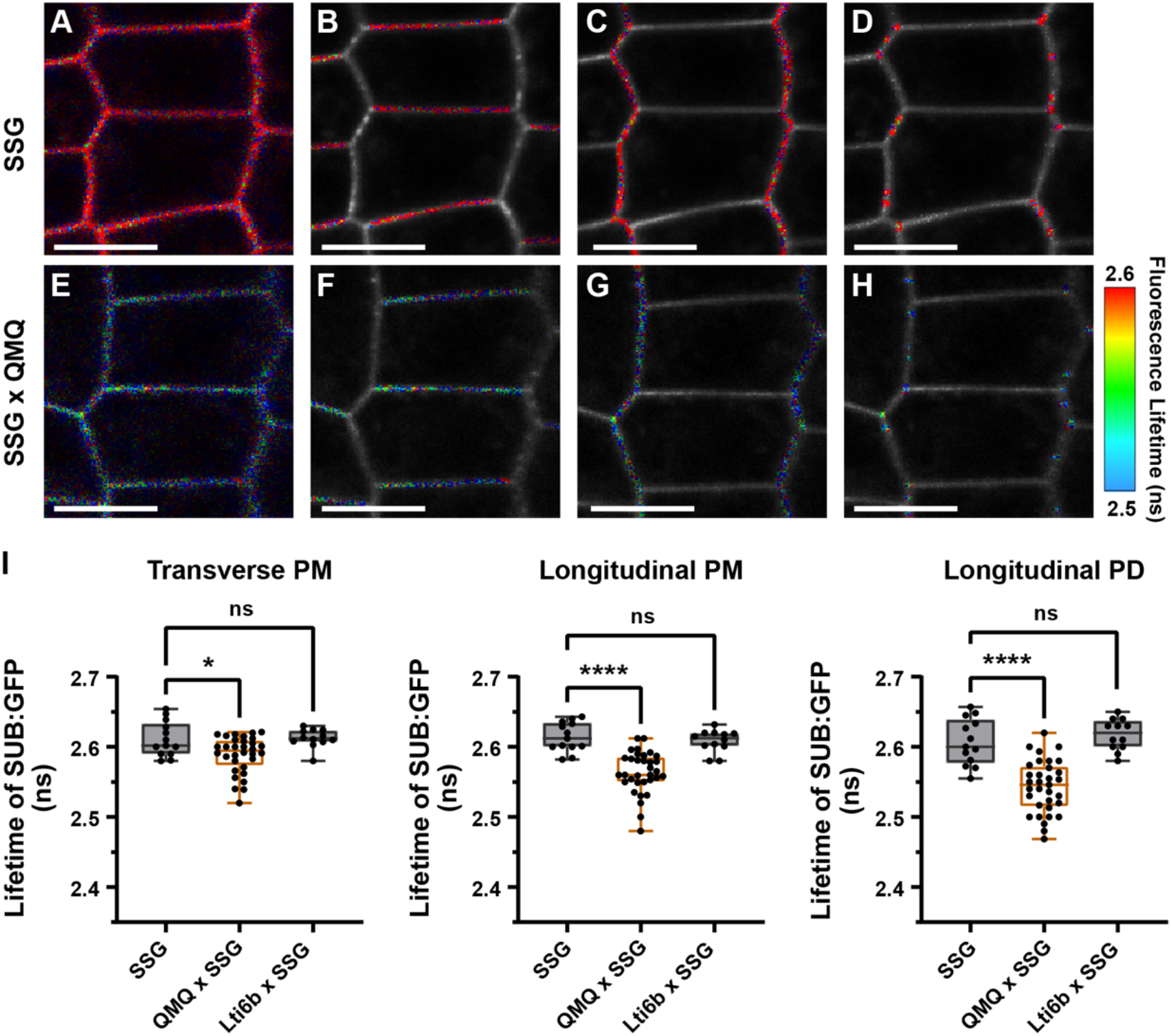
QKY physically interacts with SUB in vivo. FRET-FLIM analysis in root epidermal cells of 5-days-old seedlings of stably transgenic Arabidopsis plants. (A-D) Fluorescence lifetime images of pSUB::SUB:GFP transgenic plants. (A) Signal collected around the entire circumference of the cells. (B-D) Red areas indicate regions of interest used for signal collection at the transverse and longitudinal PM and at PD located at the longitudinal edges of the cells. (E-H) Fluorescence lifetime images of transgenic plants harboring pSUB::SUB:GFP and pQKY::mCherry:QKY. Color bar denotes the false color code for pSUB::SUB:GFP fluorescence lifetime. (I) Quantification of the pSUB::SUB:GFP fluorescence lifetimes in (A-H) at the transverse and longitudinal PM and the longitudinal PD, respectively. The transgenic plants carrying the pSUB::SUB:GFP, or pSUB::SUB:GFP and pQKY::mCherry:QKY reporters were in the *qky-9* background. Box and whisker plots are shown. Note the more pronounced reduction of pSUB::SUB:GFP lifetime along the longitudinal PM and longitudinal PD compared to the transverse PM. 12≤n≤33, each data point represents a mean of at least five cells per root from two independent transgenic lines. *P<0.02, ****P<0.0001, ns: not significant. One-way ANOVA with Tukey’s multiple comparison test. Experiments were repeated at least two times with similar results. Box and whiskers plots show the median values (middle bars) and the 25th to 75th percentiles (box). The whiskers show the smallest and largest values. Abbreviations: Lti6b, p2×35S::Lti6b:2xmCherry; QMQ, pQKY::mCherry:QKY; SSG, pSUB::gSUB:GFP. Scale bars: 8 μm.

### The C2A-B domain of QKY is required for interaction with SUB in vivo

Next, we mapped the SUB interaction domain of QKY. We first confirmed previous results (Vaddepalli *et al*, 2014) that the region comprising the C2A-D domain of QKY can interact with the intracellular domain of SUB in a Y2H assay (Fig. 5A-B). To map the interaction domain in more detail, we then generated a series of deletions within the C2A-D domain of QKY and tested their ability to interact with the intracellular domain of SUB in a Y2H assay. We found that the 187 amino acid linker region (L) separating the C2A and C2B domains was sufficient for interaction in this assay.

**Figure 5.**
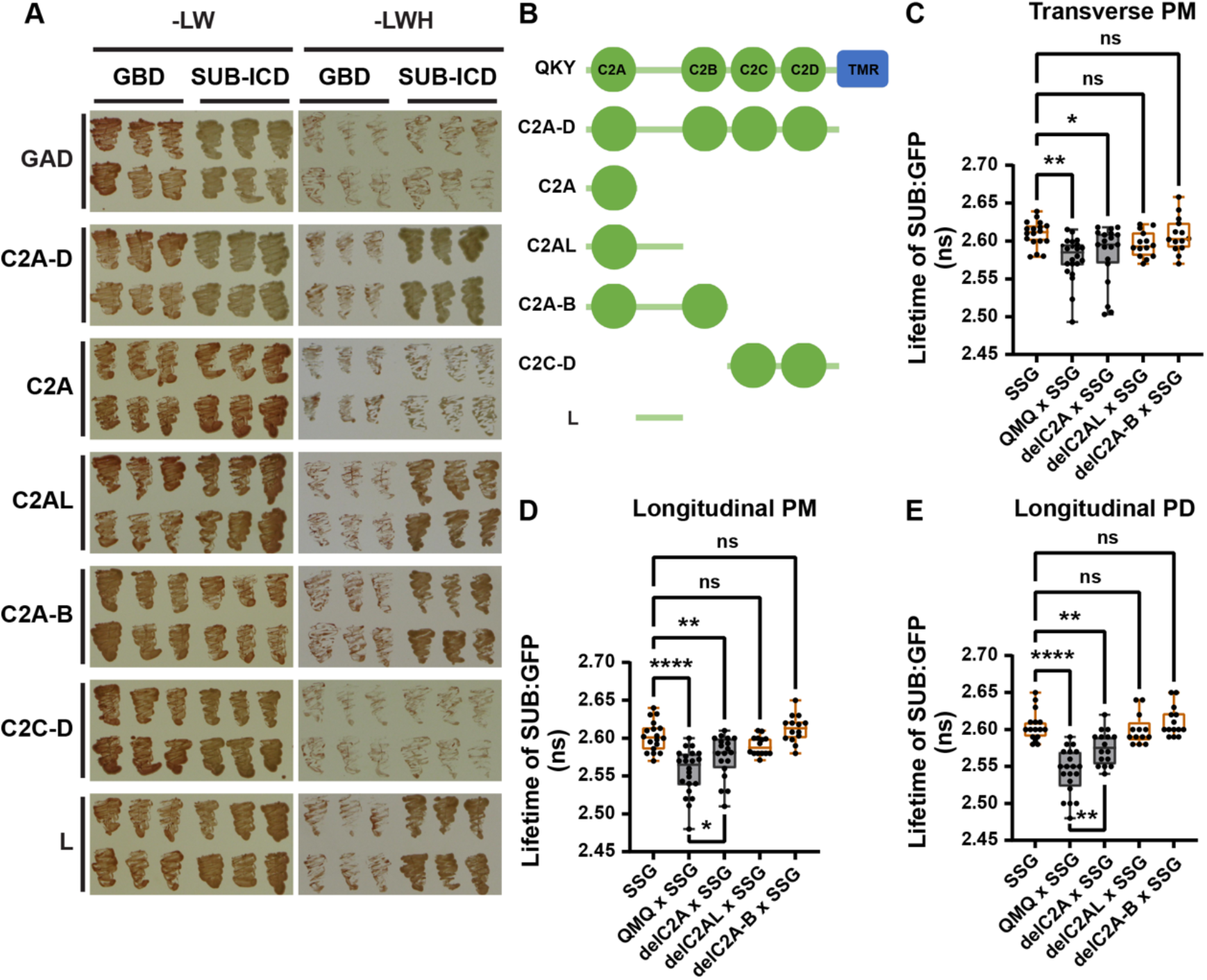
Mapping the SUB interaction region of QKY. (A) Yeast two-hybrid assay involving different truncated QKY variants fused to the GAL4 activating domain (GAD) and the SUB intracellular domain (SUB-ICD) fused to the GAL4 DNA-binding domain (GBD). Growth on -LW panel indicates successful transformation of both plasmids and on -LWH panel indicates presence or absence of interaction. (B) Cartoon depicting the different truncated QKY variants. (C-E) FRET-FLIM analysis in root epidermal cells of 5-days-old seedlings of stably transgenic Arabidopsis plants carrying pSUB::SUB:GFP combined with different mCherry:QKY variants. All the transgenic plants were in *qky-9* background. Box and whiskers plots are shown. 13≤n≤22, each data point represents a mean of at least five cells per root from two independent transgenic lines. *P<0.04, **P<0.01, ****P<0.0001, ns: not significant. One-way ANOVA with Tukey’s multiple comparison test. Experiments were repeated at least two times with similar results. Box and whiskers plots show the median values (middle bars) and the 25th to 75th percentiles (box). The whiskers show the smallest and largest values. Abbreviations: QMQ, pQKY::mCherry:QKY; SSG, pSUB::gSUB:GFP.

Next, we generated *qky-9* lines homozygous for different pQKY::mCherry:QKY variants exhibiting sequential deletions up to and including the C2D domain. We observed a PD-like localization pattern for pQKY::mCherry:QKY variants carrying a deletion of the C2A-B region. Interestingly, however, the stronger PD-related signals along the longitudinal cell periphery of pQKY::mCherry:QKY were not observed in the pQKY::mCherry:QKYdelC2A or pQKY::mCherry:QKYdelC2A-B variants (Fig. S3A-I). pQKY::mCherry:QKY variants that carried additional deletions of the C2C and C2D domains exhibited an ER-like pattern (Fig. S3J-O). The results suggest that the C2A-B domain of QKY is not required for PD localization but is involved in the preferential accumulation of QKY along the longitudinal periphery of epidermal root cells, that the C2C-D region mediates PD localization, and that the TMR is required for anchoring of QKY in the ER.

To test if the C2A-B domain was required for physical interaction with SUB in vivo we performed FRET/FLIM experiments. By crossing, we generated F1 seedlings carrying one copy of pSUB::SUB:GFP and one copy of different pQKY::mCherry:QKY variants. Compared with the mean fluorescent lifetime of pSUB::SUB:GFP in combination with pQKY::mCherry:QKY, we found a small increase in pSUB::SUB:GFP mean fluorescence lifetime when pSUB::SUB:GFP was combined with the pQKY::mCherry:QKYdelC2A construct in root epidermal cells of 5-day seedlings (Fig. 5C-E). However, its mean fluorescence lifetime was still significantly reduced compared to the pSUB::SUB:GFP control in the absence of pQKY::mCherry:QKY. When pSUB::SUB:GFP was combined with pQKY::mCherry:QKYdelC2AL and pQKY::mCherry:QKYdelC2A-B reporters, its mean fluorescence lifetime returned to the level of pSUB::SUB:GFP control in the absence of pQKY::mCherry:QKY (Fig. 5C-E). We obtained similar results for the transverse and longitudinal PMs and the longitudinal PD. Unfortunately, mapping the interaction domain of SUB is difficult to do in vivo as SUB variants carrying alterations in its ECD or intracellular domain are retained in the ER where they undergo ER-associated degradation (ERAD) (Vaddepalli *et al*, 2011; Hüttner *et al*, 2014).

### Physical interaction between QKY and SUB is required to ensure high SUB levels at the cell surface

*QKY*-mediated stabilization of SUB at the cell surface (Song *et al*, 2019; Chaudhary *et al*, 2021) (Fig. 6) counteracts the constitutive endocytosis and degradation of SUB (Gao *et al*, 2019; Song *et al*, 2019). It remains unclear, however, if physical interaction between QKY and SUB is required for this process. The pQKY::mCherry:QKY variants carrying progressive N-terminal deletions of the C2 domains failed to complement the *qky-9* phenotype (Fig. 6) (Table1, Table 2) (Fig. S6) indicating that all C2 domains are necessary for QKY function. To test whether the C2A-B SUB interaction domain controls the amount of SUB at the cell surface, we investigated the effect of the pQKY::mCherry:QKYdelC2A, pQKY::mCherry:QKYdelC2AL, and pQKY::mCherry:QKYdelC2A-B deletion constructs on the level of pSUB::SUB:GFP signal at the PM of root epidermal cells in *qky-9* seedlings. We found that all three constructs were unable to rescue pSUB::SUB:GFP levels (Fig. 6B,C). The results support the notion that the C2A-B domain is essential for maintaining high cell surface SUB levels required for the control of root hair patterning and floral morphogenesis. They also suggest that this process requires the physical interaction between QKY and SUB.

**Figure 6.**
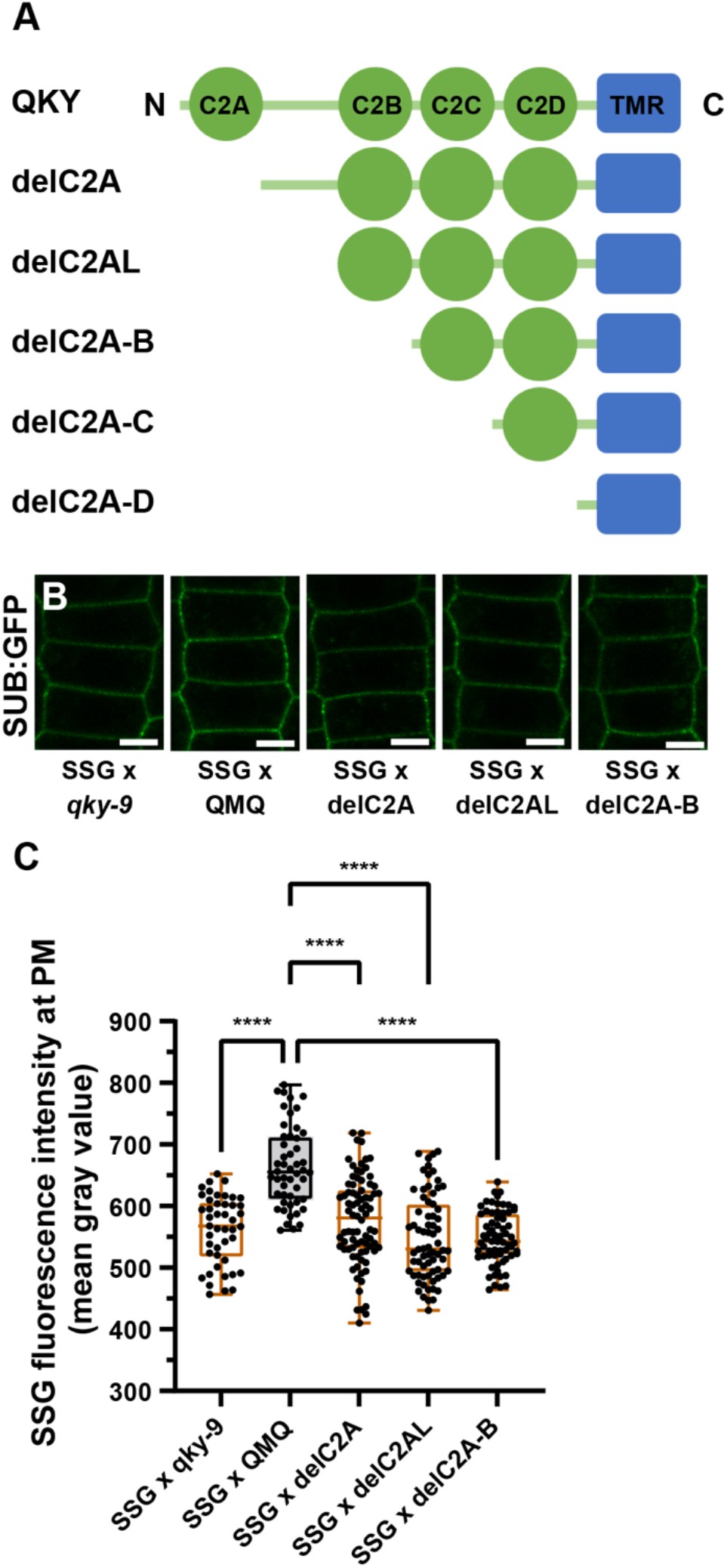
The accumulation of SUB:GFP in QKY mutants. (A) Cartoon depicting the different QKY mutants carrying progressive N-terminal deletions of the C2A-D domain. (B) Confocal micrographs depicting SUB:GFP signal intensity at the PM in root epidermal cells of 6-days-old seedlings of transgenic plants carrying combinations of pSUB::SUB:GFP with different mutant mCherry:QKY variants. All the transgenic plants were in *qky-9* background. (C) Quantification of the data in (B). Box and whiskers plots are shown. 31≤n≤70, each data point represents a mean gray value of PM in one cell across 15 different roots from two independent lines. ****P<0.0001. One-way ANOVA with Tukey’s multiple comparison test. Experiments were repeated at least two times with similar results. Box and whiskers plots show the median values (middle bars) and the 25th to 75th percentiles (box). The whiskers show the smallest and largest values. Abbreviation: SSG, pSUB::gSUB:GFP. Scale bars: 5 μm.

### SUB undergoes QKY-dependent homo-oligomerization

There is evidence that plant RKs such as BRI1 can form homo-oligomers (Russinova *et al*, 2004; Wang *et al*, 2005, 2015a). One way QKY could affect the architecture of SUB complexes at the cell surface is by controlling the oligomerization of SUB. Therefore, we investigated whether SUB undergoes homo-oligomerization. We first performed Y2H experiments with the ECD or intracellular domains of SUB. Robust yeast growth was observed on the selective medium when the ECD, but not the intracellular domain, was present as bait and prey (Fig. 7A, Fig. S7). The result demonstrates that the ECD of SUB can interact with itself in this assay. In complementary experiments we tested for interaction in root epidermal cells of 5-day seedlings expressing functional pSUB::SUB:EGFP (Vaddepalli *et al*, 2011) or pSUB::SUB:EGFP and pUBQ10::SUB:mCherry (Chaudhary *et al*, 2020) reporters. In FRET/FLIM experiments we observed a clear reduction in the mean fluorescence lifetime of pSUB::SUB:EGFP in the presence of pUBQ10::SUB:mCherry indicating that SUB forms homo-oligomers in vivo (Fig. 7B). Next, we explored if SUB oligomerization depended on QKY. To this end we measured the steady-state fluorescence anisotropy (FA) of pSUB::SUB:EGFP in root epidermal cells of 5-day wild-type and *qky-9* seedlings. We observed a significant increase in the FA value of pSUB::SUB:EGFP in *qky-9* compared with wild type (Fig. 7C). Moreover, when comparing the FA values of pSUB::SUB:GFP in *qky-9* or pSUB::SUB:GFP pQKY::mCherry:QKY *qky-9*, we found a significant decrease in pSUB::SUB:GFP FA value when pQKY::mCherry:QKY was present (Fig. 7D-L) supporting the notion that QKY promotes SUB oligomerization. We hypothesized that higher levels of SUB could lead to stronger SUB oligomerization. To test this assumption, we investigated whether the amount of SUB present at the PM affects the homo-oligomerization of SUB. To this end, we compared the FA of pSUB::SUB:EGFP in wild type and *sub-1*, a null allele of *SUB* (Chevalier *et al*, 2005). We found a modest and statistically nonsignificant increase in pSUB::SUB:EGFP FA in *sub-1*, suggesting that SUB concentration at the PM has a small, if any, effect on SUB oligomerization (Fig. 7C).

**Figure 7.**
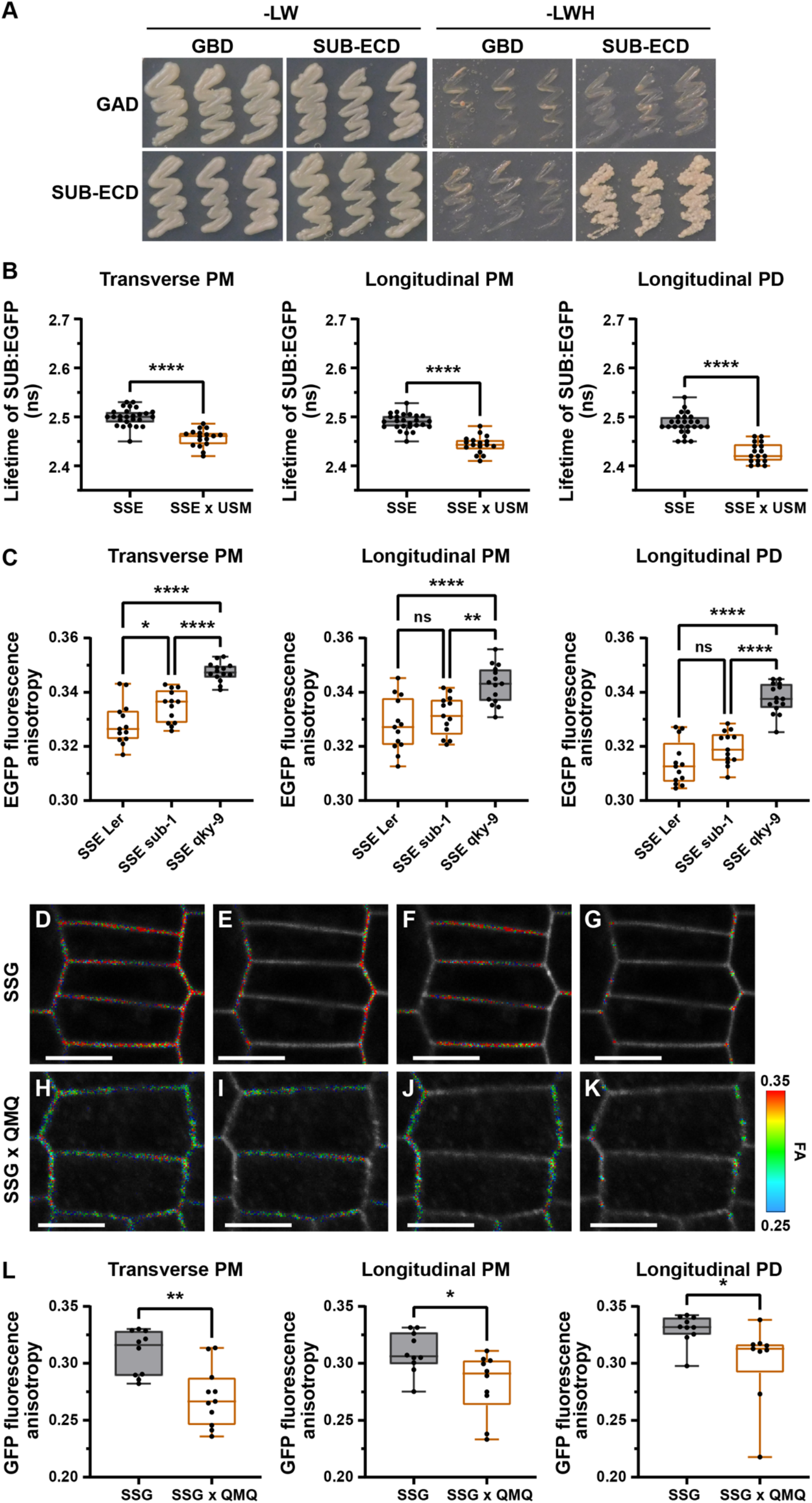
SUB undergoes QKY-dependent homo-oligomerization. (A) Yeast two-hybrid assay involving SUB extracellular domain (SUB-ECD) fused to the GAL4 activating domain (GAD) and SUB-ECD fused to the GAL4 DNA-binding domain (GBD). Growth on -LW panel indicates successful transformation of both plasmids and on -LWH panel indicates presence or absence of interaction. (B) FRET-FLIM analysis in root epidermal cells of 5-days-old seedlings of stably transgenic plants carrying pSUB::SUB:EGFP or pSUB::SUB:EGFP and pUBQ10::SUB:mCherry reporters at the transverse PM, longitudinal PM and longitudinal PD, respectively. Box and whiskers plots are shown. 17≤n≤26, each data point represents a mean of at least five cells per root. ****P<0.0001. Unpaired t test, two-tailed P-values. (C) Quantitative analysis of SUB:EGFP fluorescence anisotropy at the transverse PM, longitudinal PM and longitudinal PD, respectively, in the indicated genetic backgrounds. Box and whiskers plots are shown. 10≤n≤16, each data point represents a mean of at least five cells per root. *P<0.03, **P<0.01, ****P<0.0001, ns: not significant. One-way ANOVA with Tukey’s multiple comparison test. (D-L) Fluorescence anisotropy analysis in root epidermal cells of 5-days-old seedlings. (D-G) Images of transgenic plants carrying pSUB::SUB:GFP. Red areas indicate regions of interest used for signal collection at the transverse and longitudinal PM and at PD located at the longitudinal edges of the cells. (H-K) Images of transgenic plants carrying pSUB::SUB:GFP and pQKY::mCherry:QKY reporters. (L) Analysis of SUB:GFP fluorescence anisotropy (FA) in stably transformed *qky-9* plants harboring pSUB::SUB:GFP or pSUB::SUB:GFP combined with pQKY::mCherry:QKY reporters, respectively. pSUB::SUB:GFP fluorescence anisotropy values were assessed at the transverse PM, longitudinal PM and longitudinal PD, respectively. The lower FA values of pSUB::SUB:GFP in *qky-9* compared with pSUB::SUB:EGFP in *qky-9* are explained by weaker expression of the hemizygous pSUB::SUB:GFP reporter (see Materials and Methods). Box and whiskers plots are shown. 9≤n≤11, each data point represents a mean of at least five cells per root from two independent lines. *P<0.03, **P<0.01. Unpaired t test, two-tailed P-values. Experiments were repeated at least two times with similar results. Box and whiskers plots show the median values (middle bars) and the 25th to 75th percentiles (box). The whiskers show the smallest and largest values. Abbreviations: QMQ, pQKY::mCherry:QKY; SSE, pSUB::gSUB:EGFP; SSG, pSUB::gSUB:GFP; USM, pUBQ10::gSUB:mCherry. Scale bars: 7 μm.

## Discussion

The available data support the notion of a modular structure of QKY, in which individual domains or their combinations perform specific functions (Fig. 8A). We showed that the N-terminal C2A-B region is required for the physical interaction of QKY with the RK SUB in vivo. The combination of C2C-D and TMR domains targets QKY to PD and the TMR domain is necessary to anchor QKY to the ER, as is the case for other MCTPs (Fig. S3) (Vaddepalli *et al*, 2014; Brault *et al*, 2019; Liu *et al*, 2018a).

**Figure 8.**
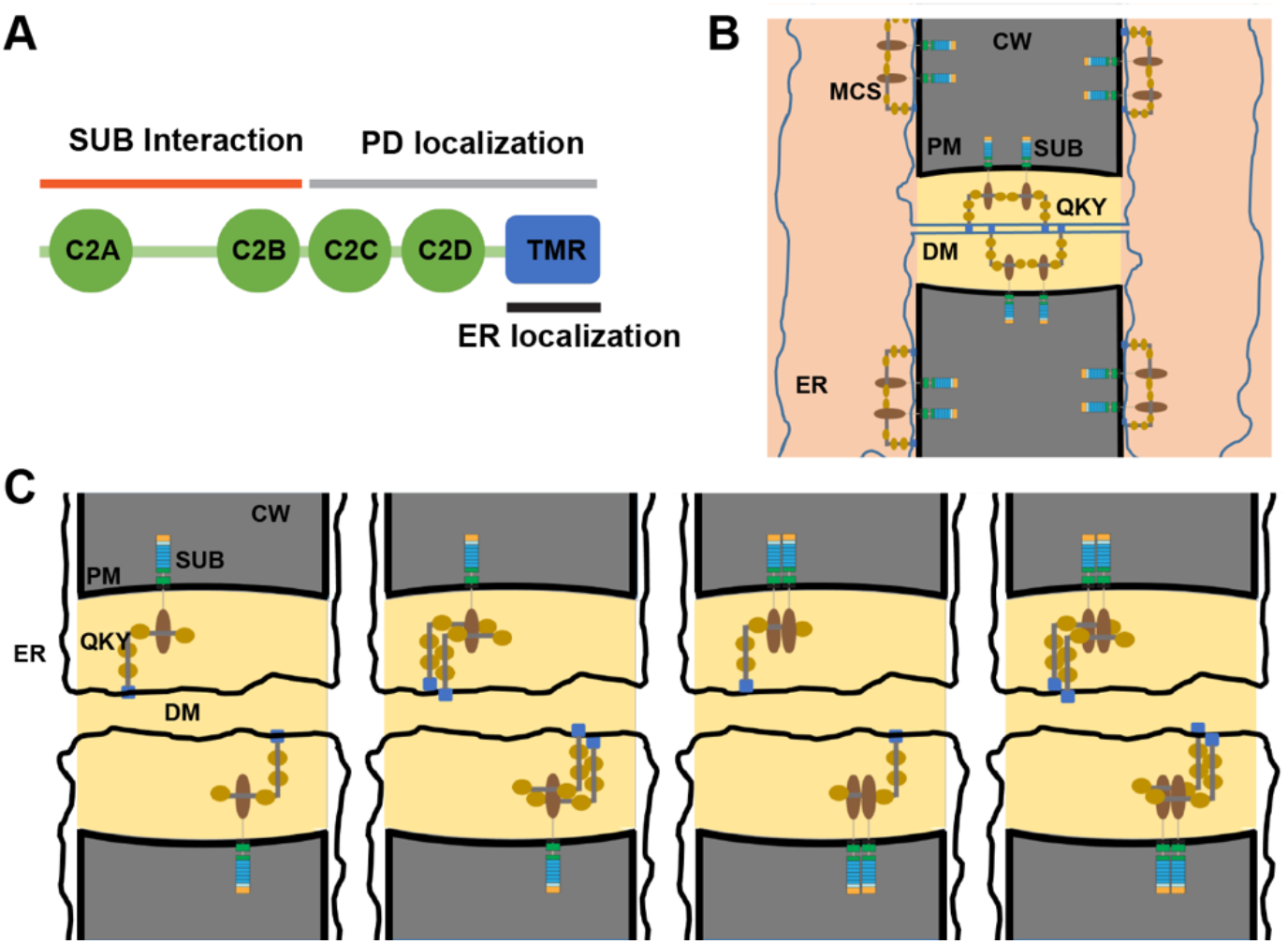
Models of QKY function. (A) Cartoon depicting the structural and functional modules of QKY. (B) Working model of SUB and QKY complexes located at PD and at sites where the cortical ER is closely associated with the PM. At both sites QKY undergoes SUB-independent homo-oligomerization. The C2A-B domain of QKY interacts with the intracellular domain of SUB thereby maintaining SUB at the PM and facilitating oligomerization of SUB. (C) Cartoon depicting the various possible complexes that could be formed between QKY and SUB. QKY and SUB could interact as monomers or interact in different oligomeric combinations. Abbreviation: CW, cell wall; DM, desmotubule; ER, endoplasmic reticulum; PM, plasma membrane; QKY, QUIRKY; SUB, STRUBBELIG; TMR, transmembrane region.

The data presented here and in previous work (Vaddepalli *et al*, 2014; Liu *et al*, 2019) reveal that reporters carrying N-terminal fusions of EGFP or mCherry to QKY driven by the endogenous *QKY* or *pUBQ10* promoters can robustly complement the phenotypes of *qky-8, qky-9, qky-14*, and *qky-17* mutants, demonstrating their functionality. Similarly, reporters containing N-terminal fusions of YFP to AtMCTP3 rescued the *mctp3 mctp4* phenotype (Brault *et al*, 2019). The collective data contrast with a recent study reporting that an N-terminal fusion of GFP to QKY driven by the endogenous promoter fails to complement the *qky-16* phenotype (Song *et al*, 2019). We do not know the reason for the discrepancy in the results, but suspect that *qky-16* is not a straightforward hypomorph but possibly a dominant-negative allele. There is a discrepancy in the literature about which domain of QKY interacts with SUB. In one study, the C2A-D domain of QKY was reported to interact with the ECD of SUB in a Y2H assay. This result in combination with proteinase K-digestions of protoplasts obtained from transgenic lines harboring tagged QKY and SUB reporters was interpreted to indicate that the C2A-D domain is extracellularly localized (Song *et al*, 2019). By contrast, we had shown earlier that the C2A-D domain of QKY interacts with the SUB intracellular domain in a Y2H assay (Vaddepalli *et al*, 2014). Moreover, we failed to detect interaction between the C2A-D domain of QKY and the ECD of SUB in multiple independent Y2H assay repetitions (Fig. S8). Finally, our two independent FRET/FLIM studies strongly suggest that the physical interaction between QKY and SUB in vivo involves the C2A-D domain of QKY and the intracellular domain of SUB, as the fluorescent tags were added to the N-terminus of QKY and the C-terminus of SUB, respectively (Fig. 4, this study) (Vaddepalli *et al*, 2014). Both studies of the in vivo QKY-SUB interaction were performed in transgenic Arabidopsis seedlings carrying functional reporters and investigated this interaction in root epidermal cells, a cell type that requires QKY and SUB function for its development (Kwak *et al*, 2005; Fulton *et al*, 2009). The combined results from our independent lines of experiments strongly suggest that the C2A-D domain of QKY localizes to the cell interior.

Our results provide deeper insight into the molecular architecture of the QKY/SUB receptor complex in vivo. Results from the Y2H analysis suggest that the N-terminal linker region of QKY flanked by the C2A and C2B domains is sufficient for the physical interaction between QKY and SUB. The FRET/FLIM data confirm a central role of the linker in the interaction in vivo. However, they also suggest that the C2A and C2B domains play an additional, albeit minor, role in this interaction, as the stepwise deletion of both domains has small but noticeable effects on the interaction. It remains to be determined whether the two domains represent additional but less important interaction interfaces or whether they form a necessary structural scaffold within QKY that allows the linker to interact with SUB. X-ray crystallography of QKY/SUB complexes will provide the answer to this question.

The available evidence strongly supports the notion that the physical interaction between QKY and SUB in Arabidopsis epidermal root cells occurs in the cytoplasm, with the C2A-B domain of QKY and the intracellular domain of SUB playing a central role in this process. The topology is consistent with the ER-localization of QKY and the PM-localization of SUB (Yadav *et al*, 2008) if one assumes that physical interaction between QKY and SUB in these cells, and possibly in other cell types, is limited to sites where the cortical ER and the PM are close enough to allow interaction (Fig. 8B). This would be the case in case of PD and is in line with our earlier model, which states that the physical interaction between QKY and SUB is confined to PD (Vaddepalli *et al*, 2014). However, this model does not explain the overall reduction in SUB at the cell periphery in *qky* mutants, which includes PD and the sections of the PM between the PD, unless one postulates that the QKY/SUB complex at the PD indirectly maintains SUB concentration at the PM between the PD. We propose a parsimonious model to explain the effect of QKY on SUB levels at the PM. We hypothesize that QKY interacts with SUB not only at PD but also at other sites along the cell periphery where the cortical ER carrying QKY is closely enough associated with the PM to allow physical interaction between QKY and SUB (Fig. 8B). In this model, physical interaction between QKY and SUB would be involved in maintaining a high level of SUB at the PM not only at PD but at numerous sites along the cell periphery.

The molecular mechanism of how QKY stabilizes SUB at the cell surface is poorly understood. Our results indicate that SUB undergoes QKY-dependent homo-oligomerization. Based on these results, we propose the following model. SUB monomers form homo-oligomers in a QKY-dependent manner at the PM and PD in epidermal root cells and possibly in other cell types. SUB homo-oligomers are less amenable to ubiquitination and internalization of SUB, a process which is inhibited by QKY (Song *et al*, 2019). For example, the ubiquitination site of SUB or its clathrin binding motif may be masked in a QKY/SUB complex. SUB oligomerization could be achieved via two non-mutually exclusive pathways. In one scenario, SUB clusters are initiated by a physical interaction between ECDs and stabilized by a subsequent physical interaction of intracellular domains with the C2A-B domain of QKY. Alternatively, an initial QKY-mediated association of the intracellular domains of the SUB may be followed by stabilization of the clusters through interactions between the ECDs. A number of questions remain. For example, different stoichiometries of the QKY/SUB complex are conceivable (Fig. 8C), and it is unclear which of these occur in vivo. It further remains to be investigated if QKY homo-oligomerization is relevant for QKY-SUB interaction and/or stabilization of SUB at PM. We will address these and other aspects in future work.

## Materials and Methods

### Plant work and genetics

*Arabidopsis thaliana* (L.) Heynh. var. Columbia (Col-0) and var. Landsberg (*erecta* mutant) (L*er*) were used as wild-type strains. Plants were grown essentially as described previously (Fulton *et al*, 2009). Plate-grown seedlings were grown in long-day conditions (16 h light/8 h dark) on half-strength Murashige and Skook (1/2 MS) agar plates supplemented with 1% sucrose. The mutant alleles *sub-1* (L*er*), *qky-9* (L*er*) and *sub-9* (Col) have been described previously (Chevalier *et al*, 2005; Fulton *et al*, 2009; Vaddepalli *et al*, 2011). The *qky-17* (L*er*) allele was generated using a CRISPR/Cas9 system in which the egg cell-specific promoter pEC1.2 controls Cas9 expression (Wang *et al*, 2015b). Two single-guide RNAs (sgRNA), sgRNA1 (5’-ACTCGGATCCTCCGCCGTCG-3’) and sgRNA2 (5’-TTACGACGAGCTCGATATCG-3) were employed. sgRNA1 binds to the region +20 to +39, while sgRNA2 binds to the region +237 to +256 of the *QKY* coding sequence. The sgRNAs were designed according to the guidelines outlined in (Xie *et al*, 2014). The *qky-17* mutant carries a frameshift mutation at position 36 relative to the *QKY* start AUG, which was verified by sequencing. The resulting predicted short QKY protein comprises 60 amino acids. The first 12 amino acids correspond to QKY, while amino acids 13-60 represent an aberrant amino acid sequence. The lines carrying pSUB::gSUB:EGFP (SSE, in *sub-1*, L*er, sub-9*), pUBQ10::gSUB:mCherry (USM, in Col), pUBQ10::EGFP:QKY (UEQ, in *sub-1*, L*er*), pUBQ10::mCherry:QKY (UMQ, in L*er*) have been previously reported (Chaudhary *et al*, 2020; Vaddepalli *et al*, 2011, 2014). The reporter lines Q4, Lti6a:GFP, Lti6b:2xmCherry, TMO7:1xGFP, and TMO7:3xGFP have been described previously (Cutler *et al*, 2000; Noack *et al*, 2022; Schlereth *et al*, 2010). The reporter constructs pGL2::GUS:GFP and pQKY::mCherry:QKY have been described earlier (Gao *et al*, 2019; Vaddepalli *et al*, 2014). Wild-type, *sub-1* and *qky-9* plants were transformed with different constructs using Agrobacterium strain GV3101/pMP90 (Koncz & Schell, 1986) and the floral dip method (Clough & Bent, 1998). Transgenic T1 plants were selected on either kanamycin (50 μg/ml) or hygromycin (20 μg/ml) plates and transferred to soil for further inspection. All the crossed materials used in this study were F1 seeds.

### Recombinant DNA work

For DNA work, standard molecular biology techniques were used. PCR fragments used for cloning were obtained using Q5 high-fidelity DNA polymerase (New England Biolabs, Frankfurt, Germany). All PCR-based constructs were sequenced. Primer sequences used in this work are listed in Table S1. The pSUB::gSUB:GFP construct was assembled using the GreenGate system (Lampropoulos *et al*, 2013). The sequences of pSUB and gSUB were amplified as previously described (Yadav *et al*, 2008; Vaddepalli *et al*, 2011). Other sequences, including GFP and the plant resistance modules, were available from the GreenGate vectors. The pSUB::gSUB:GFP was assembled into the intermediate vectors and then combined into the pGGZ0001 destination vector with a standard GreenGate reaction. The pCambia2300-based progressive N-terminal deletions of QKY carrying EGFP driven by the *UBQ10* promoter (pUBQ10::mCherry:QKYdel) were described previously (Vaddepalli *et al*, 2014). To generate the deletion constructs of QKY fused to mCherry driven by the endogenous *QKY* promoter (pQKY::mCherry:QKYdel), the fragment of pQKY::mCherry was digested with KpnI/SpeI from plasmid pQKY::mCherry:QKY, and subcloned into the KpnI/SpeI digested pUBQ::mCherry:QKYdel. For the generation of pGADT7-SUB carrying ECD or ICD, the fragment of pGBKT7-SUB carrying the ECD or ICD was digested by NdeI/XmaI and subcloned into the NdeI/XmaI-digested pGADT7. Similarly, in order to generate pGBKT7-QKYC2A-D, the fragment of pGADT7-QKYC2A-D was digested by NdeI/XmaI and subcloned into pGBKT7 digested with NdeI/XmaI. In order to generate the pGADT7 carrying the various truncated fragments of QKY, the different truncated fragments of QKY were amplified with the following primers: NdeI/QKY(-TM)_F and QKY(C2A-B)/XmaI_R for the fragment of C2A-B, QKY(C2C-D)/NdeI_F and QKY(-TM)/XmaI_R for the fragment of C2C-D, NdeI/QKY(-TM)_F and C2ALinker_XmaI_R for the fragment of C2AL, NdeI/QKY(-TM)_F and C2A_XmaI_R for the fragment of C2A, C2ALinker_NdeI_F and C2ALinker_XmaI_R for the fragment of L. Fragments were digested by NdeI/XmaI and subcloned into the NdeI/XmaI digested pGADT7. All clones were verified by sequencing. Primers are listed in Table S1.

### Yeast two-hybrid assay

The Matchmaker yeast two-hybrid system (Takara Bio Europe, Saint-Germain-en-Laye, France) was employed and experimental procedures followed the manufacturer’s recommendations. The pGBKT7 plasmids containing either the SUB extracellular (ECD) or intracellular (ICD) domain were described previously (Bai *et al*, 2013). The construct pGADT7-QKYC2A-D is equivalent to a previously described construct pGADT7-QKYΔPRT_C (Vaddepalli *et al*, 2014). In order to assess possible interactions in yeast, the different combinations of pGBKT7 and pGADT7 plasmids were co-transformed into the yeast strain AH109. Transformants were selected on synthetic complete (SC) medium lacking leucine and tryptophan (-LW) at 30°C for 3 days. To examine yeast-two-hybrid interactions, the transformants were grown on solid SC medium lacking leucine and tryptophan (SC-LW) or leucine, tryptophan and histidine (-LWH). Standard -LWH growth medium was supplemented with 2.5 mM 3-amino-1,2,4-triazole (3-AT) to minimize false positive signals. Yeast were grown for 3 days at 30°C.

### Microscopy

Floral organs and siliques were imaged using a Leica SAPO stereo microscope equipped with a digital MC 170 HD camera (Leica Microsystems GmbH, Wetzlar, Germany). Clearing and imaging of ovules was performed as reported previously (Vijayan *et al*, 2021). Ovule morphology was investigated using an Olympus BX61 upright microscope. For the analysis of root epidermal cell patterning, 6-days-old seedlings were imaged using an Olympus BX61 upright microscope equipped with an XM10 monochrome camera (Olympus Europe, Hamburg, Germany). The number of H-position and N-position cells in at least ten seedlings were scored and the relative ratio of H-position with hair and N-position without hair were deduced. For the quantitative analysis of pGL2::GUS:GFP-expressing cells in the root epiderms, 6-days-old seedlings of different genotypes carrying the pGL2::GUS:GFP reporter were counterstained with 5 μg/ml propidium iodide and examined using a FV3000 confocal laser scanning microscope (Olympus Europe, Hamburg, Germany). A high sensitivity detector (HSD) and a 60x water immersion objective (NA 1.2) were employed. Scan speed was set at 4.0 μs/pixel (image size 1024×1024 pixels), line average at 2, and the digital zoom at 1. GFP was excited using a 488 nm diode laser (2% intensity) and emission was detected at 500 to 540 nm. Propidium iodide was excited using a 561 nm diode laser (1% intensity) and emission detected at 584-653 nm. To assess pQKY::mCherry:QKY (QMQ) subcellular localization, confocal microscopy was performed on root epidermal cells of 6-days-old seedlings using an Olympus FV1000 confocal microscope and on epidermal cells of several stage 13 floral organs using an Olympus FV3000 confocal microscope. High sensitivity detectors (HSDs) and a 60x water immersion objective (NA 1.2) were employed in both microscopes. Scan speed was set at 4.0 μs/pixel (image size 1024×1024 pixels) and line average at 2. The mCherry fluorescence excitation was performed with a 561 nm diode laser (5% intensity) and detected at 584-653 nm. For the colocalization of mCherry:QKY with the ER marker Q4, 6-days-old seedling roots were imaged using an Olympus FV3000 confocal microscope equipped with HSD detectors. In some instances, the seedlings of the ER marker Q4 were counterstained with 4 μM FM4-64 (Molecular Probes) for 5 minutes. A 60x water immersion objective (NA 1.2) was employed. Scan speed was set at 2.0 μs/pixel (image size 1024×1024 pixels), line average at 2, and the digital zoom at 2. The mCherry fluorescence and the FM4-64 stain were excited using a 561 nm diode laser (7% intensity for mCherry and 2% intensity for FM4-64) and emission was detected at 584-653 nm. GFP was excited using a 488 nm diode laser (1% intensity) and emission was detected at 500-540 nm. To observe the subcellular localization of pQKY::mCherry:QKY mutant variants, the root epidermal cells of 6-days-old seedlings were imaged using an Olympus FV3000 confocal microscope equipped with a HSD and a 60x water immersion objective (NA1.2). Scan speed was set at 4.0 μs/pixel (image size 1024×1024 pixels) and the digital zoom at 4. The mCherry fluorescence excitation was performed with a 561 nm diode laser (7% intensity) and detected at 584-653 nm. Images were adjusted for color and contrast using ImageJ/Fiji software (Schindelin *et al*, 2012). For the quantification of SUB-GFP accumulation at the PM, epidermal cells of root meristems of 6-days-old seedlings in the corresponding plant line were imaged using an Olympus FV3000 microscope equipped with a HSD and a 60x water immersion objective (NA 1.2) at digital zoom 4. Scan speed was at 4.0 μs/pixel (image size 1024×1024 pixels) and line average at 2. GFP was excited using a 488 nm diode laser (1% intensity) and emission was detected at 500 to 540 nm. For the direct comparisons of fluorescence intensities, laser, pinhole, and gain settings of the confocal microscope were kept identical when capturing the images in different QKY mutant backgrounds. The mean gray values of GFP fluorescence signal intensity at the PM were measured in 5-10 cells from each image by creating a region of interest covering the PM area using ImageJ/Fiji.

### FRET-FLIM and fluorescence anisotropy measurements

FRET-FLIM was performed using an Olympus FV3000 microscope equipped with a time-correlated single photon counting (TCSPC) device (LSM upgrade kit, PicoQuant, Berlin, Germany). Root epidermal cells of 5-days-old seedlings co-expressing SUB:GFP and mCherry:QKY were imaged with a 60x water immersion objective (NA 1.2) at digital zoom 4. Scan speed was 4.0 μs/pixel (image size 512×512 pixels). The FLIM filter cube DIC560 for GFP/RFP was employed. GFP fluorescence lifetimes were measured with two photon-counting PMA hybrid 40 detectors and a pulsed 485 nm diode laser (LDH-D-C-485, PicoQuant) using a laser pulse rate of 40 MHz. For each image, a minimum of 300 photons per pixel were acquired with a TCSPC resolution of 25.0 ps (image size 512×512 pixels). Results were analyzed using SymPhoTime 64 software 2.7 (PicoQuant) using n-exponential reconvolution and an internally calculated instrument response function (IRF). The lifetime fitting model parameter as 1 was defined (n=1). The analysis results with a correctional factor (χ^2^) between 0.9 and 1.9 were accepted. Intensity-weighted average lifetime (τ Av Int) of membrane ROIs was taken as the final value for each FLIM image. Steady-state fluorescence anisotropy was performed and analyzed as reported previously (Chaudhary *et al*, 2021; Chaudhary & Schneitz, 2022). Note that if one has to work with weak reporter expression and therefore around the minimum number of detected photons required for a meaningful result (on average 15 photons per pixel of a ROI detected in the channel belonging to the perpendicular detector of the PicoQuant system (channel 1 in the SymPhoTime 64 software 2.7), maximum scan time of 5 min), the actual calculated anisotropy value is also influenced to some extent by the number of photons collected. In such a scenario, a lower number of photons leads to a low FA value, and a higher number of photons leads to a higher FA value.

### Statistical analysis

Statistical analysis was performed with PRISM 9.4.0 software (GraphPad Software, San Diego, CA, USA). All statistical tests and P-values are described in the respective figure legends.

## Acknowledgements

We thank members of the Schneitz lab for helpful comments and discussion. We also thank Farhah Assaad (Technical University of Munich), Ziqiang Patrick Li and Emmanuelle Bayer (CNRS-University of Bordeaux) for the Q4, Lti6a:GFP and Lti6b:2xmCherry reporters. We thank Ajeet Chaudhary for help with the yeast two-hybrid method, and acknowledge Rachele Tofanelli and Tejasvinee Mody for their help with ovule imaging. We further thank Ajeet Chaudhary and Sebastian Wolf for suggestions and comments. We also acknowledge support by the Center for Advanced Light Microscopy (CALM) of the TUM School of Life Sciences.

## Supplement

**Figure S1:**
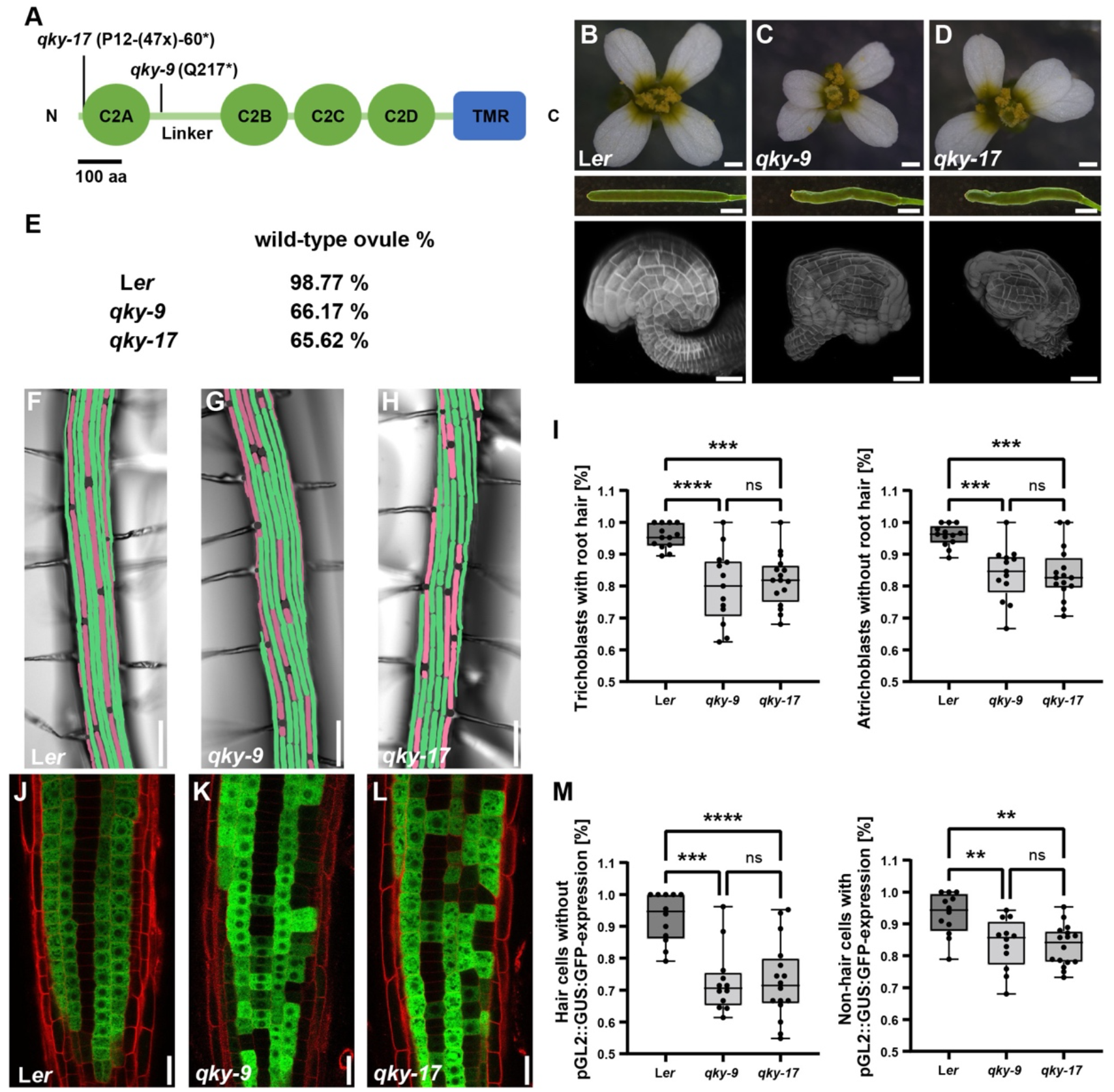
Phenotypic comparison of L*er, qky-9* and *qky-17*. (A) Cartoon depicting the structure of QKY protein. The four C2 domains in the N-terminal and the transmembrane region (TMR) in C-terminal are indicated. The positions of the *qky-9* and *qky-17* mutations are also shown. The *qky-9* allele causes a stop codon at position 217 just following the C2A domain. The putative null allele *qky-17* was generated using a CRISPR/Cas9 system and predicted to encode a short protein comprising 60 amino acids. The first 12 amino acids correspond to QKY, while amino acid 13-60 represent an aberrant amino acid sequence. (B-D) Floral organ morphology of L*er, qky-9, qky-17*. Upper panel: stage 13 flower, middle panel: stage 15 silique, bottom panel: stage 3-V ovule. (E) The proportion of ovules with wild-type appearance in L*er, qky-9* and *qky-17*, respectively. For each genotype, 7 to 10 different carpels from at least two different plants were analysed (430≤n≤458). (F-H) Micrographs showed epidermal root hair cell patterning in 7-days-old seedlings of L*er* (F), *qky-9* (G) and *qky-17* (H). (I) Quantification of results shown in (F-H). The percentage of H position hair cells with root hairs (left panel) and that of N position hair cells without root hairs (right panel) are shown, respectively. Box and whisker plots are shown. 13≤n≤16, each data point represents the percentage in per different root, which was calculated by at least 15 cells of each cell type. ***P<0.001, ****P<0.0001, ns: not significant. One-way ANOVA with Tukey’s multiple comparison test. Experiments were repeated at least two times with similar results. Box and whiskers plots show the median values (middle bars) and the 25th to 75th percentiles (box). The whiskers show the smallest and largest values. (J-L) Expression pattern of the pGL2::GUS:GFP reporter in L*er* (J), *qky-9* (K) and *qky-17* (L). (M) Quantification of the results shown in (J-L). The percentage of H position hair cells without expressing the pGL2::GUS:GFP reporter (left panel) and that of N position hair cells expressing the pGL2::GUS:GFP reporter (right panel) are shown, respectively. Box and whisker plots are shown. 12≤n≤16, each data point represents the percentage in per different root, which was calculated by at least 15 cells of each cell type. **P<0.01, ***P<0.001, ****P<0.0001, ns: not significant. One-way ANOVA with Tukey’s multiple comparison test. Experiments were repeated at least two times with similar results. Box and whiskers plots show the median values (middle bars) and the 25th to 75th percentiles (box). The whiskers show the smallest and largest values. Note that *qky-9* and the putative null allele *qky-17* show identical phenotypes, indicating that the C2A domain of QKY potentially present in cells of *qky-9* does not have residual function or act in a dominant-negative fashion. Thus, *qky-9* represents a putative null allele as well. Scale bars: (B-D) flowers: 400 μm, siliques: 2mm, ovules: 20 μm. (F-H) 100 μm, (J-L) 20 μm.

**Figure S2:**
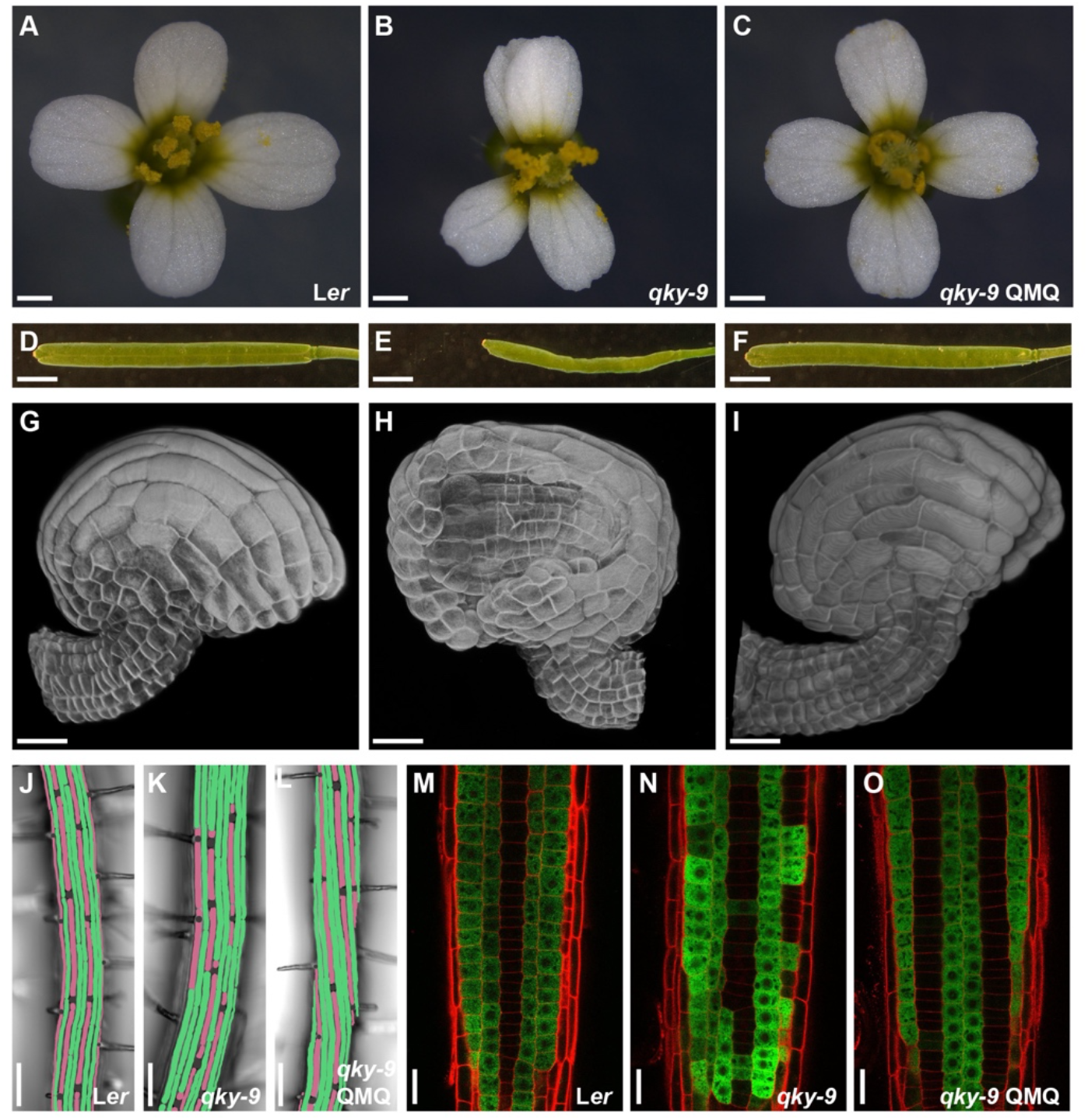
pQKY::mCherry:QKY rescues the *qky-9* phenotype. pQKY::mCherry:QKY (QMQ) encodes a functional fusion protein. (A-C) Stage 13 flower morphology of L*er* (A), *qky-9* (B) and QMQ *qky-9* (C). (D-F) Silique shape (stage 15) of L*er* (D), *qky-9* (E) and QMQ *qky-9* (F). (G-I) Stage 3-V ovule morphology of L*er* (G), *qky-9* (H) and QMQ *qky-9* (I). (J-L) Micrographs showed epidermal root hair cell patterning in 7-days-old seedlings of L*er* (J), *qky-9* (K) and QMQ *qky-9* (L). (M-O) Expression pattern of the pGL2::GUS:GFP reporter in L*er* (M), *qky-9* (N) and QMQ *qky-9* (O). Confocal micrographs show optical sections of epidermal cells of root meristems of 7-days-old seedlings. The cell wall was counterstained with propidium iodide. Scale bars: (A-C) 0.5 mm, (D-F) 2 mm, (G-I) 20 μm, (J-L) 100 μm, (M-O) 20 μm.

**Figure S3:**
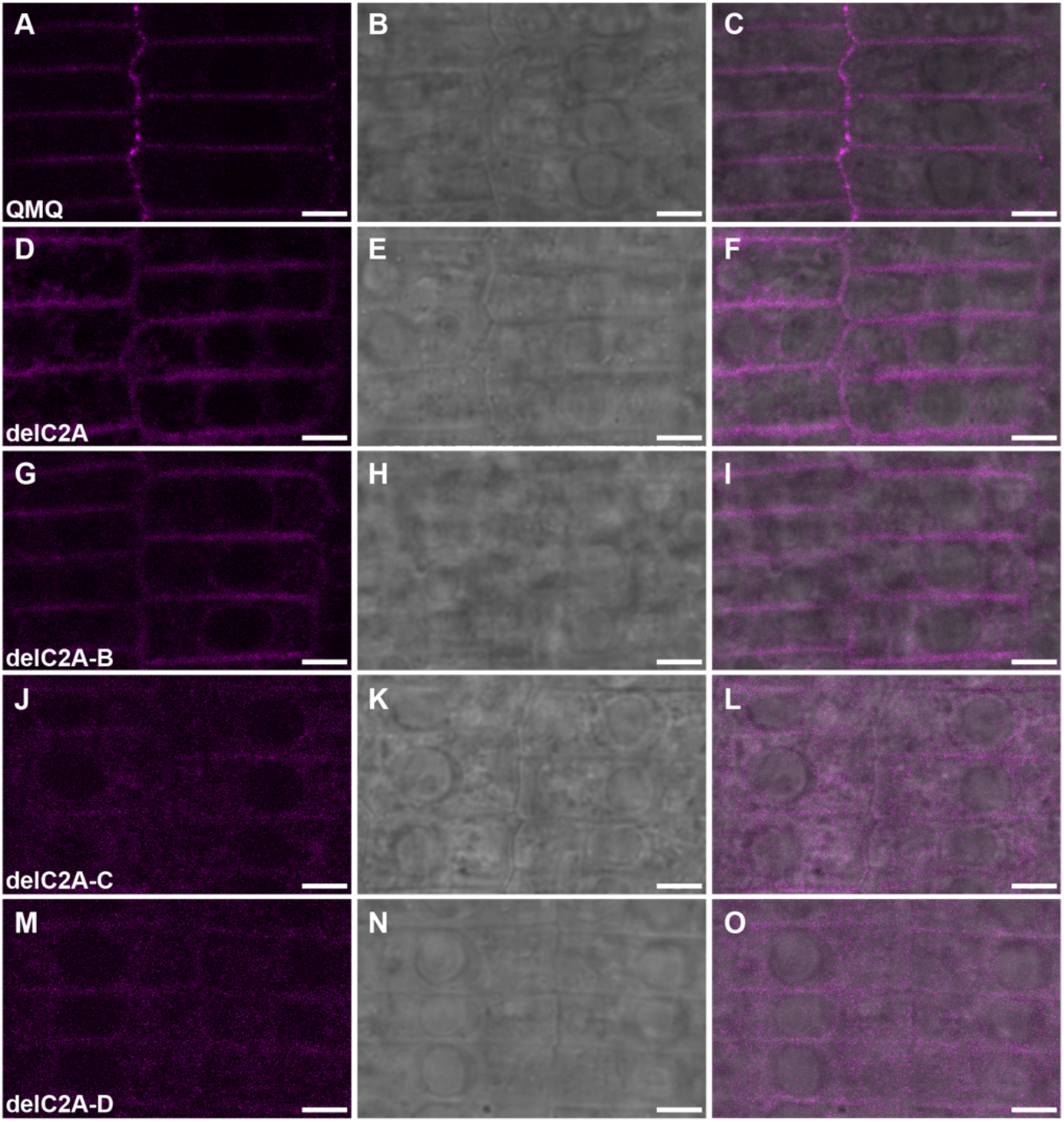
Subcellular localization of different pQKY::mCherry:QKYmut reporters. Confocal micrographs of root epidermal cells of 6-days-old seedings of pQKY::mCherry:QKY (QMQ) mutant in *qky-9* background. Left panels:mCherry signal. Middle panels: DIC channel. Right panel: merge. (A-C) Note the punctate pattern at the PM in QMQ *qky-9*. (D-I) The punctate pattern was attenuated, but clear PM-like localization is still observable in QMQdelC2A *qky-9* and QMQdelC2A-B *qky-9*. (J-O) Reduced PM-like localization and more diffuse cytoplasmic signal distribution in QMQdelC2A-C *qky-9* and QMQdelC2A-D *qky-9*. Scale bars: 5 μm.

**Figure S4:**
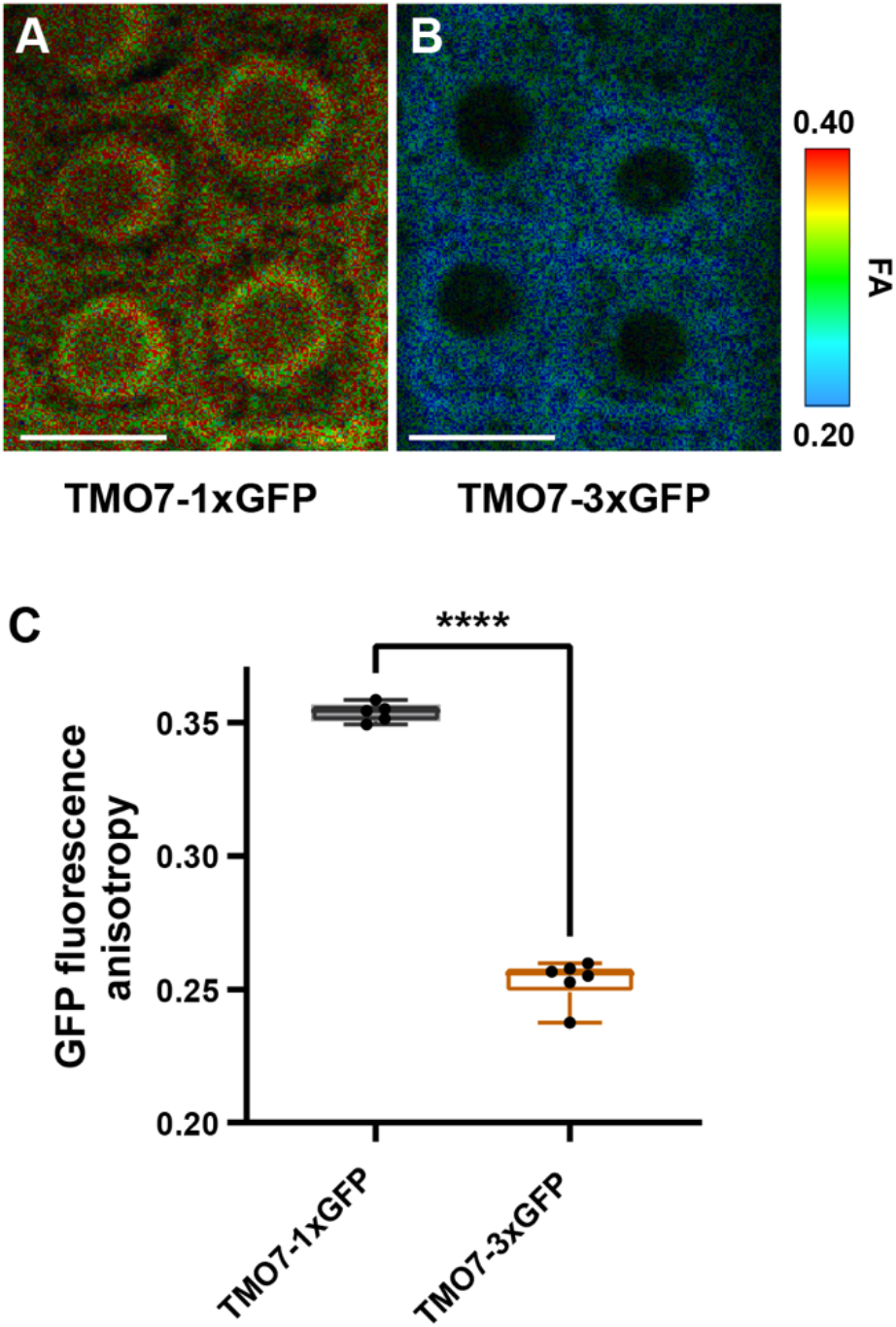
Fluorescence anisotropy of TMO7:1xGFP and TMO7:3xGFP in root epidermal cells. (A-B) Images of root epidermal cells of 5-days-old seedlings carrying TMO7:1xGFP and TMO7:3xGFP, respectively. Color bar denotes the false color code for SUB:GFP fluorescence anisotropy. (C) Quantification of GFP florescence anisotropy in (A-B). Note the significant reduction of anisotropy in TMO7:3xGFP. Box and whisker plots are shown. 5≤n≤6, each data point represents a mean of at least five cells per root. ****P<0.0001. Unpaired t test, two-tailed P-values. Experiments were repeated at least two times with similar results. Box and whiskers plots show the median values (middle bars) and the 25th to 75th percentiles (box). The whiskers show the smallest and largest values. Scale bars: 9 μm.

**Figure S5:**
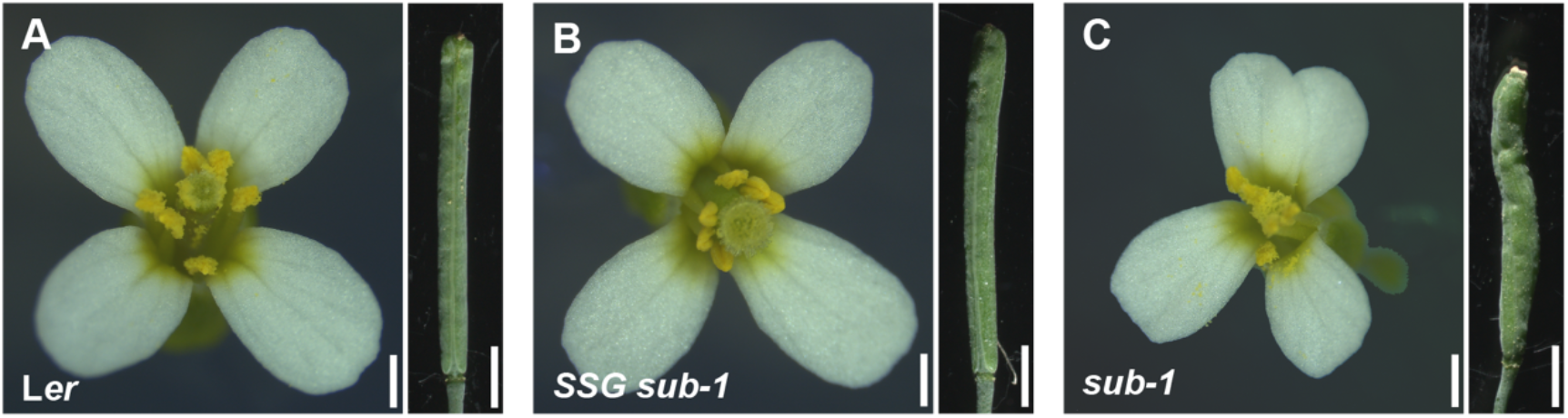
Functionality of the pSUB::SUB:GFP (SSG) reporter. Micrographs of stage 13-14 flowers and siliques are shown. The genotypes are indicated. Note the wild-type appearance of *SSG sub-1* flowers (compare A with B and C). Scale bars: flowers, 0.5 mm; siliques, 2 mm.

**Figure S6:**
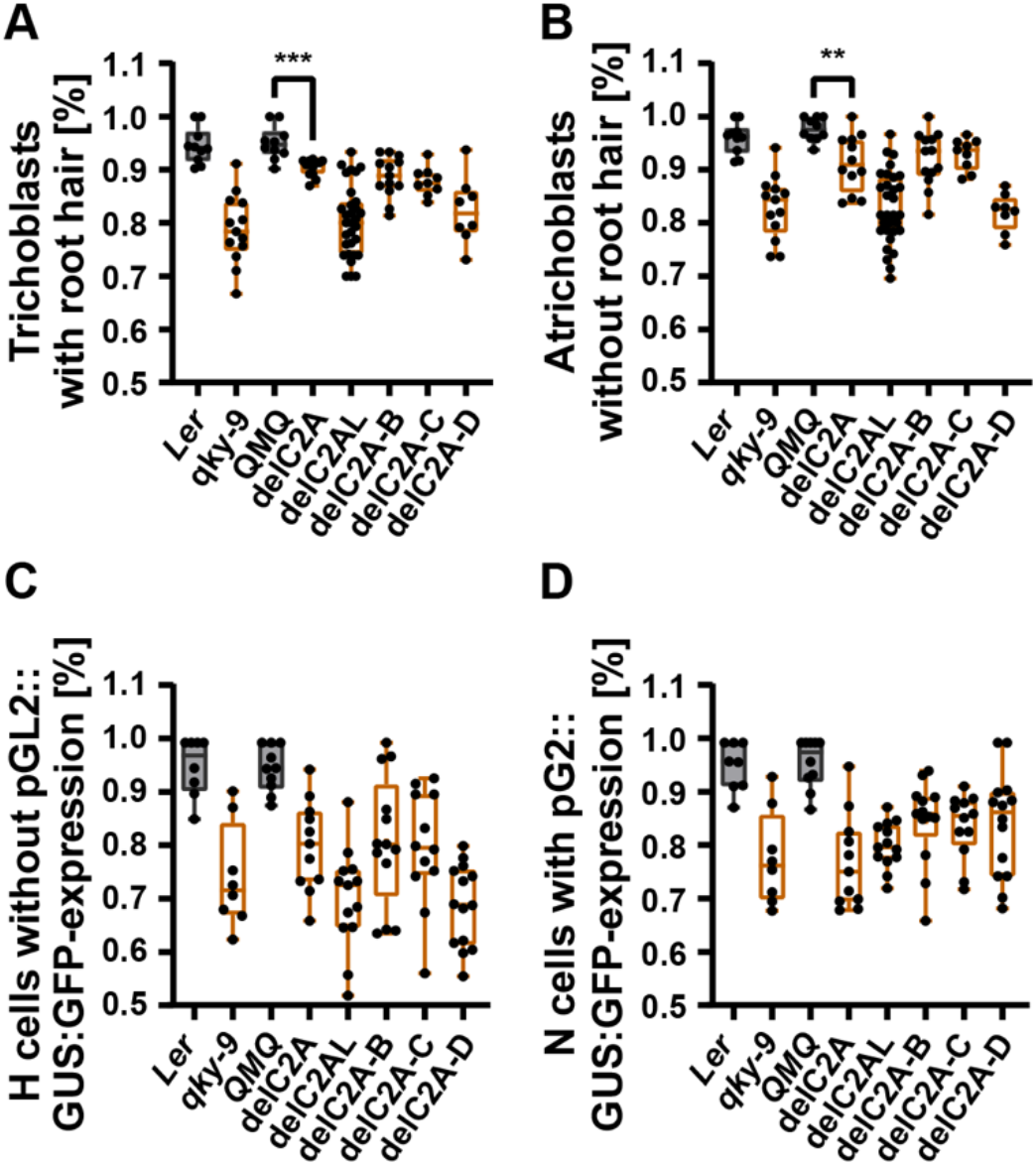
Specification of root epidermal cell types in *qky-9* mutants carrying progressive N-terminal deletions of the C2A-C2D domain. All tested progressive N-terminal deletions within the C2A-C2D domain affect QKY-dependent root hair patterning. (A-B) The percentage of H position hair cells with root hair and that of N position hair cells without root hair in different genotypes, respectively. Box and whisker plots are shown. 8≤n≤31, each data point represents the percentage in per root from two independent lines, which was calculated by at least 15 cells of each cell type. **P<0.01, ***P<0.001. Black boxes represent no significant differences in the data between L*er* and pQKY::mCherry:QKY (QMQ). Orange boxes represent significant differences in the data of *qky-9* and QKY mutants compared to L*er* and QMQ. Unpaired t test, two-tailed P-values. Experiments were repeated at least two times with similar results. Box and whiskers plots show the median values (middle bars) and the 25th to 75th percentiles (box). The whiskers show the smallest and largest values. (C-D) The percentage of H position hair cells without expressing the pGL2::GUS:GFP reporter and that of N position hair cells expressing pGL2::GUS:GFP reporter in different genotypes, respectively. Box and whisker plots are shown. 8≤n≤14, each data point represents the percentage in per root from two independent lines, which was calculated by at least 15 cells of each cell type. Black boxes represent no significant differences in the data between L*er* and QMQ. Orange boxes represent significant differences in the data of *qky-9* and QKY mutants compared to L*er* and QMQ. Unpaired t test, two-tailed P-values. Experiments were repeated at least two times with similar results. Box and whiskers plots show the median values (middle bars) and the 25th to 75th percentiles (box). The whiskers show the smallest and largest values.

**Figure S7:**
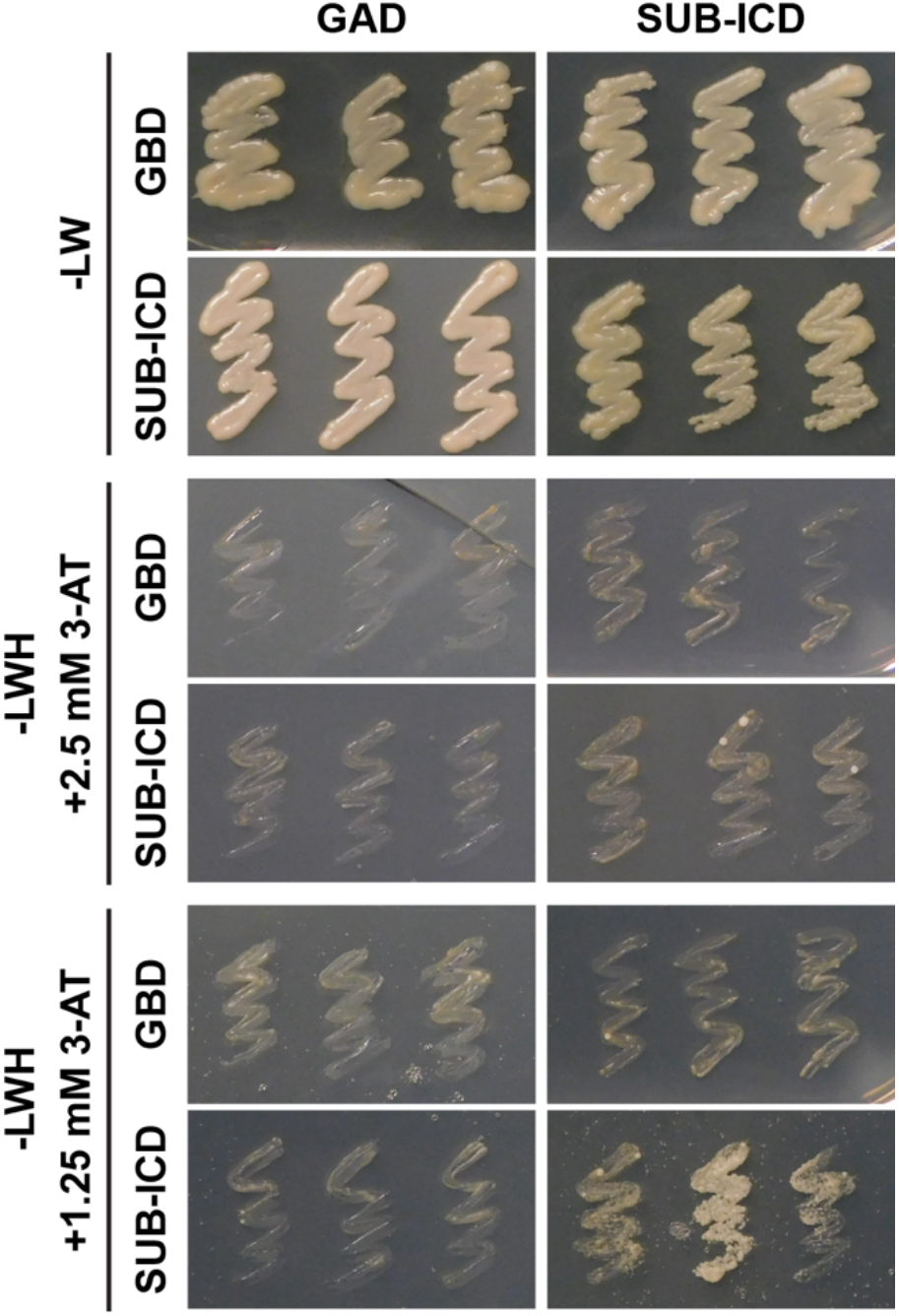
The intracellular domain (ICD) of SUB does not robustly interact with itself in a Y2H assay. Either standard conditions (2.5 mM 3-AT) or conditions with reduced stringency (1.25 mM 3-AT) were applied. Note the weak and variable growth of yeast colonies carrying BD-ICD and AD-ICD grown in the presence of 1.25 mM 3-AT. First colonies were detected after 5 days. Yeast growth was stopped after 7 days. The experiment was repeated twice with identical results.

**Figure S8:**
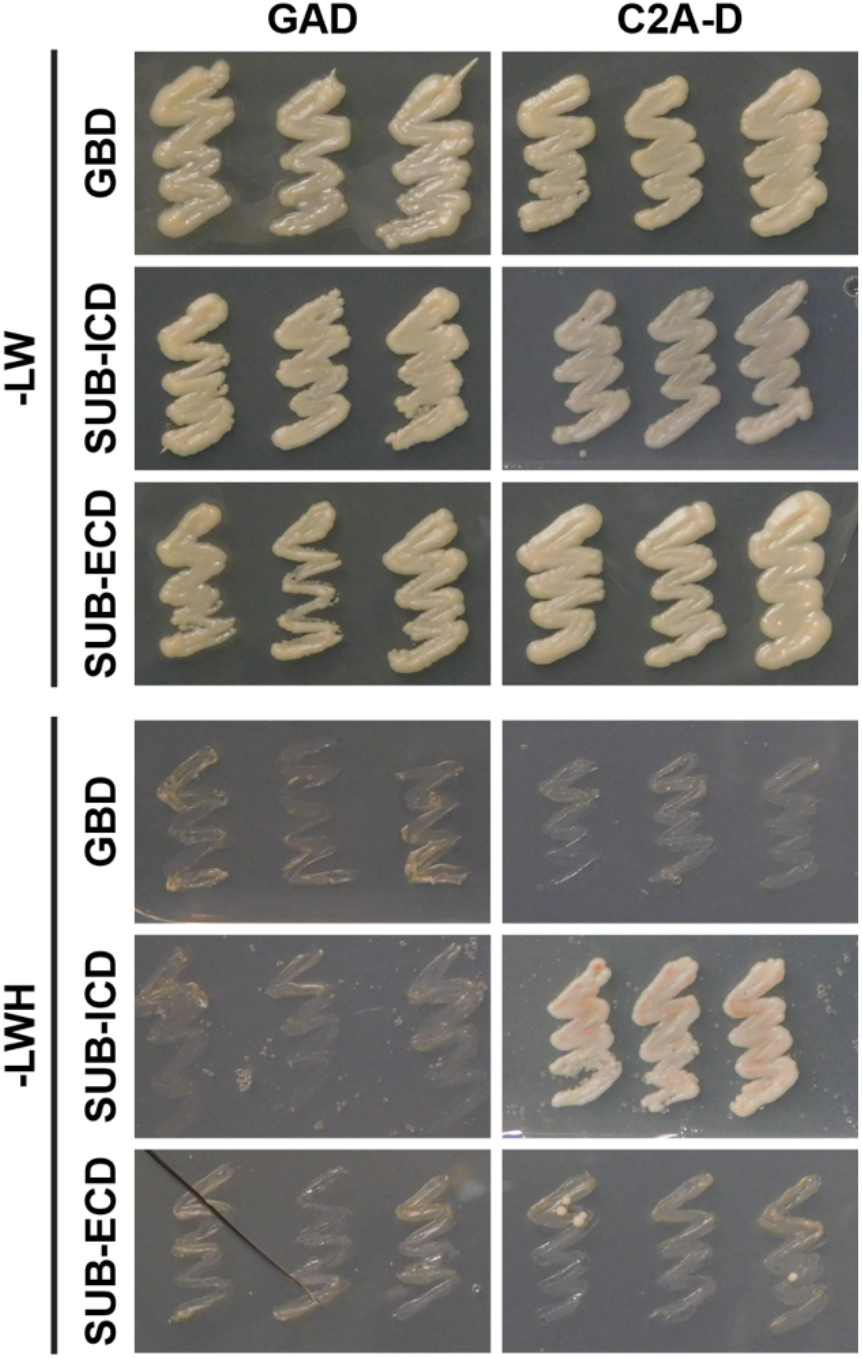
The ECD of SUB does not interact with the C2A-D domain of QKY in a Y2H assay. The intracellular domain (ICD) of SUB serves as a positive control. Standard conditions (2.5 mM 3-AT) were applied. Yeast growth was stopped after 3 days. Identical results were obtained when stopping yeast growth after 7 days. The experiment was repeated twice with identical results.

**Table S1:**
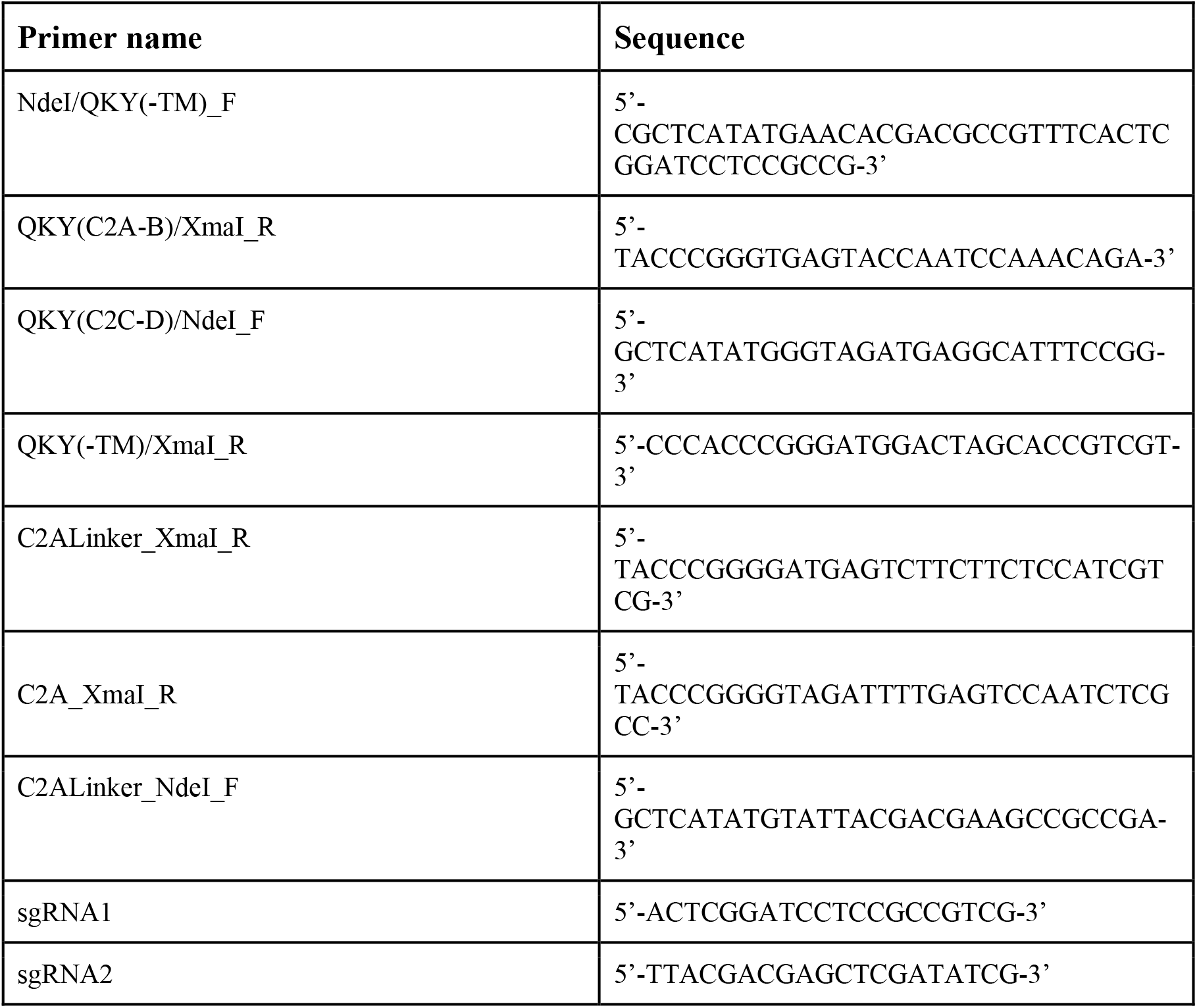
Primers used in this study.

